# Vesicle Navigation of Microtubule Ends Distinguished by A Single Rate-Constant Model

**DOI:** 10.1101/2020.07.21.214023

**Authors:** M.W. Gramlich, S. Balseiro Gómez, S. M. Ali Tabei, M. Parkes, S. Yogev

## Abstract

Axonal motor driven cargo utilizes the microtubule cytoskeleton in order to direct cargo, such as presynaptic vesicle precursors, to where they are needed. This transport requires vesicles to travel up to microns in distance. It has recently been observed that finite microtubule lengths can act as roadblocks inhibiting vesicles and increasing the time required for transport. Vesicles reach the end of a microtubule and pause until they can navigate to a neighboring microtubule in order to continue transport. The mechanism by which axonal vesicles navigate the end of a microtubule in order to continue mobility is unknown. In this manuscript we model experimentally observed vesicle pausing at microtubule ends in *C. elegans*. We show that a single rate-constant model reproduces the time vesicles pause at MT-ends. This model is based on the time a vesicle must detach from its current microtubule and re-attach to a neighboring microtubule. We show that vesicle pause times are different for anterograde and retrograde motion, suggesting that vesicles utilize different proteins at plus and minus end sites. Last, we show that vesicles do not likely utilize a tug-of-war like mechanism and reverse direction in order to navigate microtubule ends.

## 1. Introduction

The length of neuronal axons necessitates a dedicated transport system to deliver cargo from the cell body over large distances. Molecular motors dynein ^1^, and kinesin superfamily members ^2^, use the uniform plus-end-out microtubule (MT) array of axons as a substrate on which to transport axonal cargo retrogradely or anterogradely, respectively. Axonal transport is crucial for neuronal viability, and its dysfunction is a hallmark of neurodegenerative diseases ^3^.

Paradoxically, despite the need to cover long-distances, cargo motility in axons is interspersed with frequent pauses ^4,5^. The fraction of time cargo spends being immotile varies considerably among cargo types and experimental systems ^6,7^. Explanations for these stalls include a tug-of-war between dynein and kinesin motors ^8,9^, interactions with physical barriers (such as other organelles, MAPs or actin) ^5,10^, and pauses at MT ends.

Tug-of-war mechanisms have been extensively studied in vitro ^9^, and in vivo ^8^. Protein barriers have been likewise studied in vitro ^11^, and in vivo ^12^. Cargo pauses at MT-ends along the axon have been less well studied with in vitro studies of structural MT-defects have focused exclusively on single MTs ^13^. Our recent in vivo experiments of synaptic vesicle precursors have shown a significant inhibitory effect at MT-ends ^7^. We focus here on understanding the recently observed in vivo experiment mechanism by which cargo navigate MT-ends ^7,14^.

All axons harbor parallel tracks of individual MT polymers, which are much shorter than the axon itself ^15–17,7,18,19^. When a cargo reaches a MT-end it must somehow detach and re-attach somewhere else in order to navigate the MT-end. This raises the question of how the cargo negotiate the transfer from one polymer to the next at MT ends. Our previous work revealed that in *C.elegans* motor neurons, most pauses during the transport of synaptic-vesicle precursors (SVPs) occur at MT tips, indicating that negotiating the transfer from one polymer to the next is rate-limiting for efficient transport in this system ^7^.

Several possible scenarios may occur at microtubule ends: (i) cargo-motor complexes may fall off at the MT-end, then the cargo diffuses until it binds a new motor somewhere else along the axon and become motile again; (ii) multiple motors bound to the cargo complexes may compete at a MT end, and the resumption of cargo motility would depend on either motor dissociation from the paused motor; (iii) the cargo-motor complex detaches from the MT at the end and the same motor-cargo reattaches to another MT along the bundle.

Studies in vitro and in non-neuronal cultures have partially addressed these possibilities. For example, dynein motors show tenacious binding of MT-minus-end at the end of their runs in vitro ^20,21^. Conversely, kinesins display a more variable behavior at the plus-end, ranging from tight binding to falling off ^22,14,23^, which depends on the specific type of kinesin examined, the presence of co-factors, the GTP-state of the MT and the experimental system. However, analyzing similar events in neurons faces two significant technical obstacles: (1) The difficulty of visualizing individual motors limits the analysis to the motility of the cargo and (2) The high density of MTs in axons does not allow to resolve which polymer a given cargo is associated with at the light-microscopy level. Hence, it is presently unknown how cargo negotiates microtubule ends to resume motility in neurons. One approach that could bridge the gap between in vitro observations and studies in neurons is modeling.

Here we develop a simple probabilistic model for the behavior of motor-vesicle complexes at MT ends during axonal transport. Importantly, the model enables us to describe the observed pausing behavior using a single parameter. We then test the model against *in vivo* live imaging of SVP transport in *C.elegans* motor neurons, where we have shown that pauses during transport occur at MT ends. We find that the model reliably explains the behavior of vesicle at MT ends. These studies provide insight into long-range transport of synaptic cargo in axons.

## 2. Experimental Methods

Raw data collection was performed as previously described ^7^. Briefly, young-adult hermaphrodite *C.elegans* expressing the synaptic vesicle precursor marker RAB-3 in the DA9 motor neuron (wyIs251[Pmig-13::GFP::RAB-3]) were paralyzed in 0.3 mM Levamisole in M9. Once paralyzed, worms were carefully transferred to M9 solution on a 10% agarose pad for imaging. This results in an effective Levamisole concentration which is significantly lower than the concentration that was suggested to affect axonal transport ^24^. Worms were maintained on the pad for no more than 20 min, although we confirmed that viability was maintained even after 4 hrs.

Fluorescence imaging was performed using a Nikon 60x CFI plan Apo VC, NA 1.4 objective on a Nikon Ti-E microscope equipped with Yokogawa CSU-X1 scan-head and a Hamamatsu C9100-50 EM-CCD camera at a frame rate of 110ms/frame, and 240 nm per pixel.

Post-analysis fluorescence movies were corrected and analyzed in imageJ. Animal movement was corrected using FIJI plugin StackReg. Kymographs were generated with KymoBuider. Intensity was averaged +/− 5 pixels transverse to the kymograph line.

## 3. Results

### 3.1. Quantifying Pause times at Microtubule ends

We first characterized MT-end locations along an axon in order to determine pause-times (Fig. 1). Cargo exhibited the ability to both traverse (blue arrow in Fig. 1 (B)) and pause (orange arrow in Fig. 1 (B)) at the same x-axis location along a kymograph. In order to determine locations of MT-ends at x-positions, we identified positions where cargo transport frequently paused, as our previous work identified these locations as microtubule ends. We required *at least* 10 cargo observed to traverse and *at least* 5 cargo were observed to pause. The combined number of traverse and pause cargo would give an uncertainty in measuring any given pause-time of < 10% for each MT-end measured (See Appendix A1 for methods).

**Figure 1:**
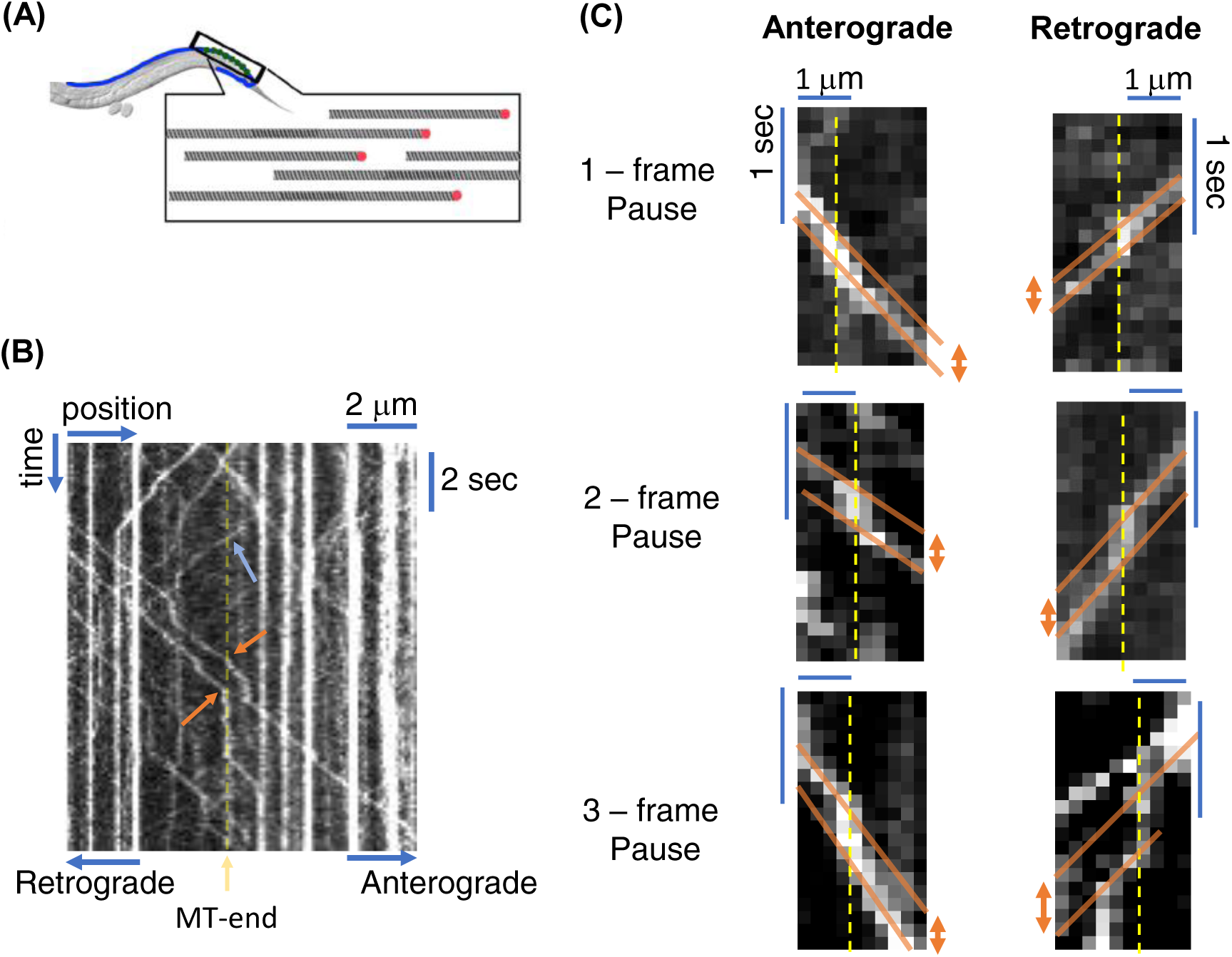
MT-end kymograph analysis method. (A) Cartoon geometry of *C. elegans* axon and microtubule network. The axon has multiple tracks of microtubules aligned parallel to each other with breaks between individual microtubules. (B) Kymograph (time/space) representation of cargo navigating an axon. A microtuble end is designated as dashed-yellow line. Vesicles are marked as traversing (blue arrow) or pausing (orange arrow) at the end. Vesicles engage in both anterograde and retrograde motion. (C) Examples of quantification of short-time vesicle pausing at microtubule-ends (dashed yellow lines). The pause-time at a MT-end is determined as the time difference between velocity traces (orange solid lines). Both anterograde and retrograde motion are quantified the same and exhibit the same pausing behavior.

We noted that some locations also had cargo stalled before imaging began and remained at the same location for the entire experiment. experiment. Although such locations likely correspond to MT ends^7^, we wished to eliminate a possible effect of cargo aggregation ^25^. Our focus in this work was on the role of MT-ends and not stalled vesicles. Any motile vesicle, which paused at the same location as a stalled vesicle, may have paused because of the stalled vesicle and not the MT-end. The imaging resolution could not distinguish between a pause caused by a MT-end and a stalled vesicle. Thus, we did not include locations that had cargo paused before imaging began.

We quantified cargo pausing at MT-ends based on velocity and time-shifts in kymograph data (Fig. 1 (C)). We particularly focused on short-time pauses because they represented the majority fraction of all cargo pausing. We distinguished between pause times by following protocol: (i) we calculated the slope of cargo trajectory before (solid orange line in all figures) reaching a known MT-end location (dashed yellow line in all figures); (ii) we then calculated the slope of cargo trajectory after the known MT-end location (solid orange line in all figures); (iii) we then determined the intercept for each trajectory either before (for anterograde motion) or after (for retrograde motion) at a fixed position (slopes are drawn to the edge of the kymograph where they are compared in Fig. 1 (C)); (iv) finally, the difference between intercepts is then the pause time for the cargo. Using this method we can easily distinguish between short-time (1-3 frames) pauses as shown by the examples in Fig 1 (C).

We note that as control we also chose random locations along the x-axis where at least 20 cargo observed to traverse but no requirement for the number of cargo pauses at that location. We used the same quantification method to determine short-time pauses at these random locations. We chose the same number of locations as MT-end locations. Last, we followed the same data aggregation method for these random locations in order to determine pause-time distribution (See Appendix A1).

### 3.2. Probability of traversing a MT-end

We first quantified the fraction of attempts that motor driven cargo walk past a MT-end. If a vesicle is on a MT that ends, then it must detach and re-attach to another MT along the axon. However, in vivo fluorescence microscopy experiments cannot distinguish which MT a vesicle is on. Thus, if a vesicle is travelling along the axon, fluorescence experiments measure the probability that it is on the MT that ends. This probability is simply the inverse of the number of laterally aligned MTs. This probability is true regardless of the motor or direction of travel. Thus, we quantified the probability that vesicles *walk past* a MT-end in order to determine if simple probabilities dictate motor pausing. It is important to note that all MTs along an axon have their polarities aligned with plus-end distal and minus-end proximal oriented ^25,17,26^.

Experimentally we observed that both retrograde (Dynein) and anterograde (kinesin-3/UNC-104) driven motion exhibit similar probability of traversing MT-ends. Retrograde driven vesicles traversed MT-end locations the majority of times observed (0.76 +/− .02). Anterograde driven motion also traversed the same MT-end locations the majority of times observed (0.71 +/− 0.02). Although anterograde traversal amount was slightly less than retrograde, it was not statistically significantly different (P = 0.594, two-tailed Students t-test). The overall fraction of vesicle traversals suggest that vesicles have approximately 1 in 4 probability of being on a MT with a MT-end, which is consistent with the range of MT tracks in the DA9 axon, as previously determined by EM reconstructions ^7^.

We controlled for random pausing effects by comparing traverse fractions at MT-ends to traverse fractions at randomly chosen locations along the axon (Fig. 2). If vesicle traversing at MT-ends is unrelated to MT-end locations, then vesicles would exhibit the same fraction of traversals at randomly chosen locations. However, there is a statistically significantly higher rate of vesicle traversals at random locations. Both retrograde driven motion (MT-end to random location; P = 5.06E-5; two-tailed t-Test) and anterograde driven motion (MT-end to random location; P = 9.24E-6; two-tailed t-Test) exhibit the same higher traversals (Retrograde: 0.92 +/− 0.02; Anterograde: 0.89 +/− 0.02; p = 0.1146; two-tailed t-Test).

**Figure 2:**
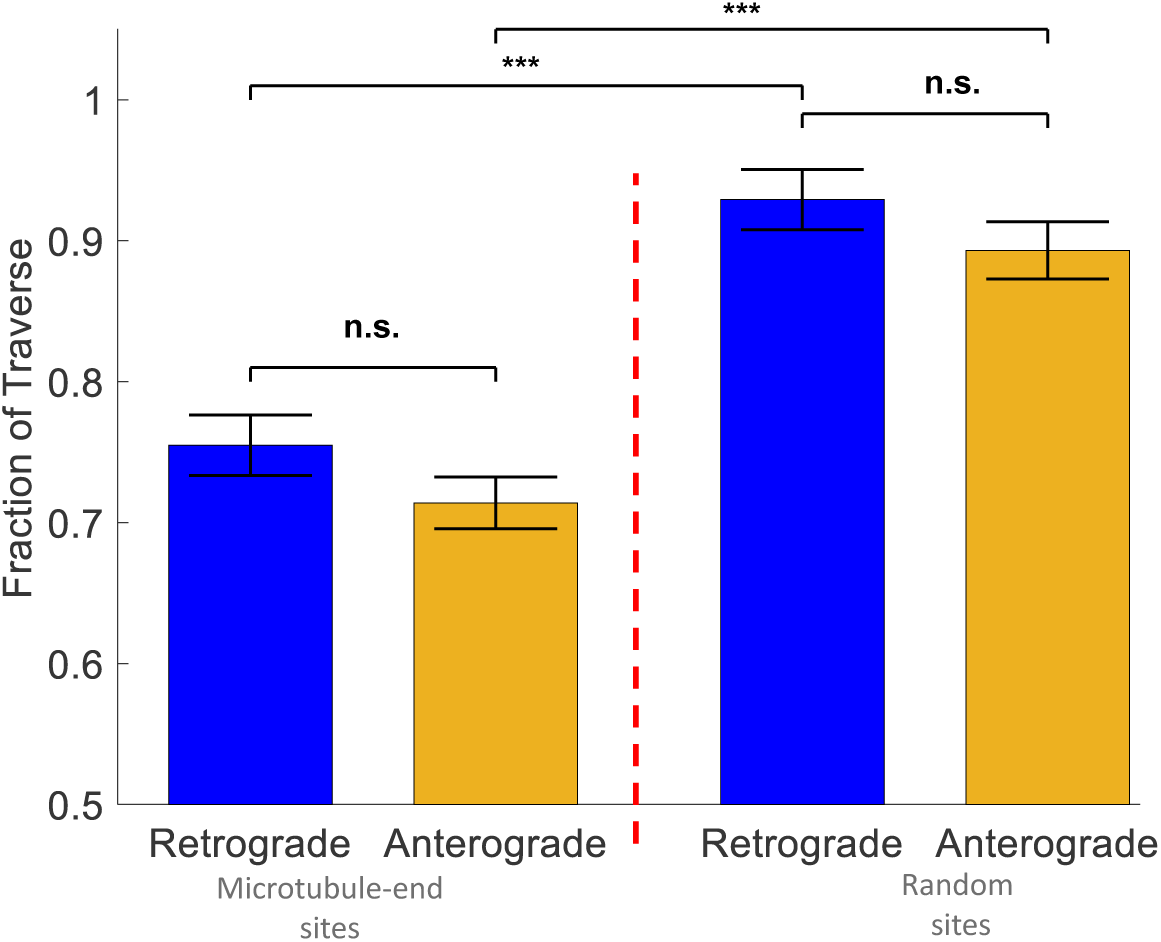
Probability Vesicles Traverse A MT-end. Both retrograde (Blue Bars) and anterograde (Yellow Bars) driven vesicles traverse MT-ends for the majority fraction of attempts (>0.7 for both). Retrograde and anterograde motion traverse a greater amount at randomly chosen axon locations, as compared to MT-end locations. *** = p < 0.01; Error-bars are determined by standard-deviations in fraction of observed traversals per location.

### 3.3. Vesicle Reversals suggest minimal tug-of-war motion

We next turned to distinguishing the dominant mechanism by which motors navigate MT-ends. There have been multiple proposed models axonal cargo use to navigate the cytoskeleton such as: (i) the “bucket-brigade” model ^27^, which suggests motors are constantly binding and detaching from vesicles but directed motion continues as if a single motor drives mobility; (ii) multiple bound motors per vesicle coordinate motion ^28,29^; (iii) a “tug-of-war” model between anterograde/retrograde motors where vesicles will occasionally engage both motors simultaneously to navigate obstructions ^9,30,4^. Distinguishing between different models is essential to understand the specific mechanisms by which vesicles navigate MT-ends.

We first focused on whether the “tug-of-war” mechanism was involved in vesicle motion by quantifying vesicle reversals. Tug-of-war was first observed in vesicles that reverse direction at random locations along the axon. This observed tug-of-war behavior was distinguished from the same motor reversing direction due to oppositely polarized MTs because axons have all MTs oriented with the same polarity ^25,17,26^. We observe that vesicles engage in reversals at MT-end locations (Fig. 3A). Indeed, both anterograde and retrograde motion exhibit reversals at MT-ends. *If vesicles utilize the “tug-of-war” mechanism then vesicle reversals at MT-ends would be a significant fraction of overall observed motion. Alternatively, if “tug-of-war” does not significantly contribute how vesicles navigate MT-ends then reversals would be a small fraction of vesicle motion.*

**Figure 3:**
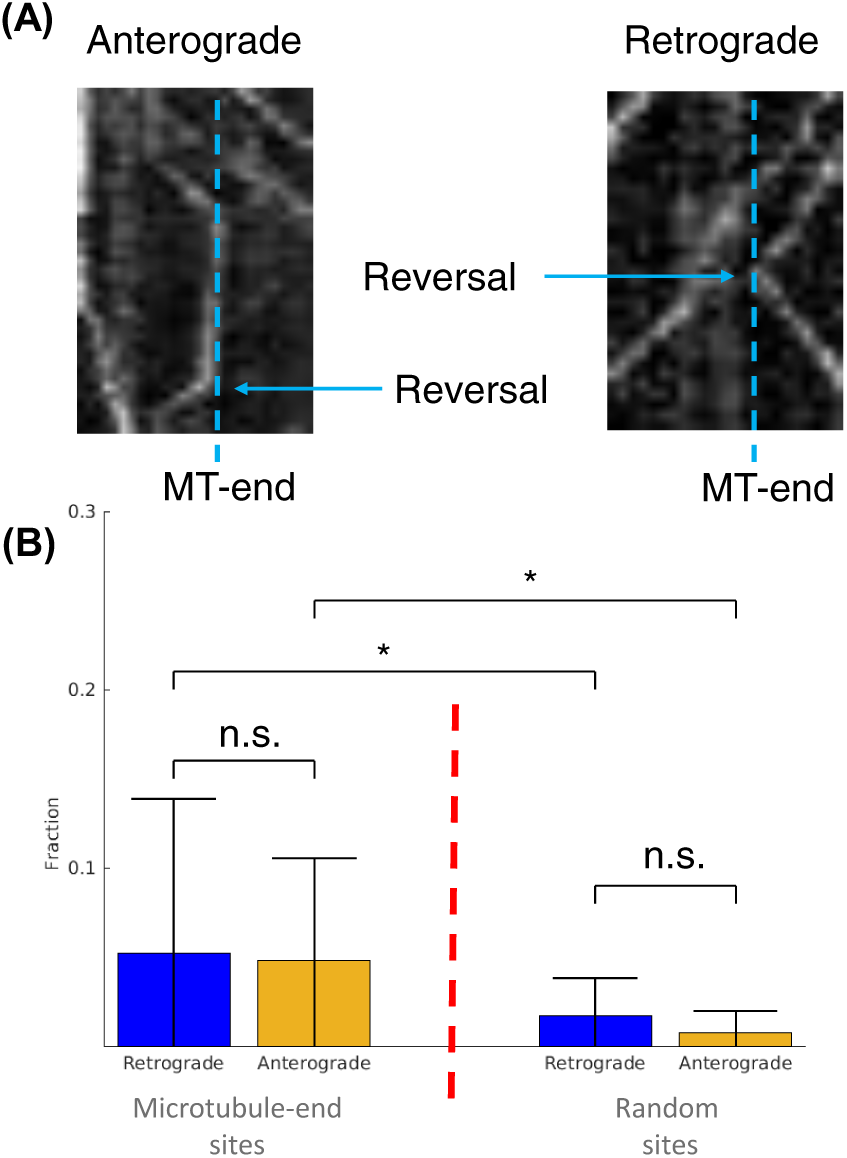
Vesicle Reversal Fraction *after* Pausing at a Microtubule-end. (A) Both anterograde and retrograde motion exhibit reversals at a microtubule-end location. Reversals are not observed to correlate with pause time. (B) The overall fraction of reversals at microtubule-ends is <0.05 for both retrograde and anterograde motion. There is no statistical difference for direction of motion at MT-end. The overall fraction of reversals at random locations along the axon is <0.02 for both retrograde and anterograde motion. There is no statistical difference for direction of motion at random locations. Error-bars are determined by standard-deviations in fraction of observed traversals per location. * = p <0.05, two-tailed student t-Test.

We observed that vesicle reversal is a small portion of overall vesicle mobility at MT-ends (Fig. 3B). Importantly, reversals are not observed to correlate with pause time (Data not shown). We determined aggregate vesicle reversals by counting reversals across 10 MT-ends in 3 different experiments and divided by the total number of tracks for the same MT-ends. Both Retrograde and anterograde motion exhibited the same overall low reversal fraction with no statistically significant difference (Retrograde = 0.05 +/−0.086, Anterograde = 0.048 +/− 0.057; P = 0.6255 from 2-tailed t-test). Thus, reversal and any potential motor transition at MT-ends is a low fraction of overall vesicle mobility at MT-ends.

We compared the fraction of observed vesicle reversals at MT-ends to observed vesicle reversals at random locations along the same axons. We determined aggregate vesicle reversals by counting reversals across 12-13 locations in 4 different experiments and divided by the total number of tracks for the same locations. Reversals at random positions along the axon occurred in less than 2% of all observed pauses for both retrograde and anterograde motion (Retrograde = 0.017 +/−0.02, N = 353; Anterograde = 0.008 +/− 0.01, N = 393; P = 0.182 from 2-tailed t-test). There was a small but significant increase in reversals at MT-ends compared to random locations for both retrograde (P = 0.0463 from 2-tailed t-test) and anterograde (P = 0.0223 from 2-tailed t-test) motion. However, the overall scale of reversals regardless of location is small compared to the total number of observed vesicles.

The small fraction of reversals suggests that *the “tug-of-war” mechanism is minimally utilized by vesicles to navigate MT-ends*.

### 3.4. Single Rate-constant Model of Cargo Navigation of MT-ends

The low-reversal results lead to the following question: *what mechanism(s) determine how long a vesicle will pause at a MT-end before resuming mobility?* If vesicles do not reverse, then what ever motor was driving the vesicle before the MT-end may likely dominate the time required for a vesicle to navigate the MT-end.

We propose the following hypothesized model/mechanism: vesicle navigation of MT-ends depends on the direction currently used by the vesicle when it encounters a MT-end. This hypothesis suggests that the time paused at MT-ends depends on the rate-constants of the motor driving the vesicle before it reaches the MT-end and any possible end-binding protein or the structure of the MT end itself that mediates vesicle binding/un-binding at the MT-end. Note that these rate-constants are not the attachment/detachment rates that are observed for single motors along a MT in vitro (See discussion section). Further, this hypothesis suggests that anterograde and retrograde motion should exhibit different distributions of pause-times at MT-ends.

We now propose a model and method to distinguish our hypothesized mechanism by which vesicles navigate MT-ends. This model is based on two observations/assumptions: (i) The probability that a vesicle will pause is only dependent on the microtubule track it is currently attached (based on the traverse fraction in Fig. 2); (ii) The time a vesicles pause depends entirely on the detachment/attachment rate-constants of the motor driving the cargo when it reaches a MT-end. With these two observation-based assumptions we created a probabilistic model and computational simulation similar to observed vesicle transport.

It is important to note that we only compare the times our simulated cargo spend at MT-ends to observed experimental times. However, we model transport along the entire axon to ensure that our model is also consistent with observed experimental kymograph results.

#### 3.4.1. Axonal Bundle Modelling

We modeled the axon as a bundle of aligned microtubules tracks ^17^, as observed experimentally (Fig. 1A). The tracks are a two-dimensional arrays with rows representing tracks of individual microtubules and columns representing a coarse-grained lattice site (Fig. 4A). Microtubules are aligned along tracks equidistant from each other in a radial fashion similar to a neuronal axon or dendrite ^31^. We set the minimum spatial resolution limit of our model to the size of a microtubule end, equal to the diameter of a single microtubule ∼24 nm (represented by boxes in Fig. 4A) ^32,33^. Single microtubules of finite-length populate these tracks with MT-ends represented by a single missing lattice site (Fig. 2A).

**Figure 4:**
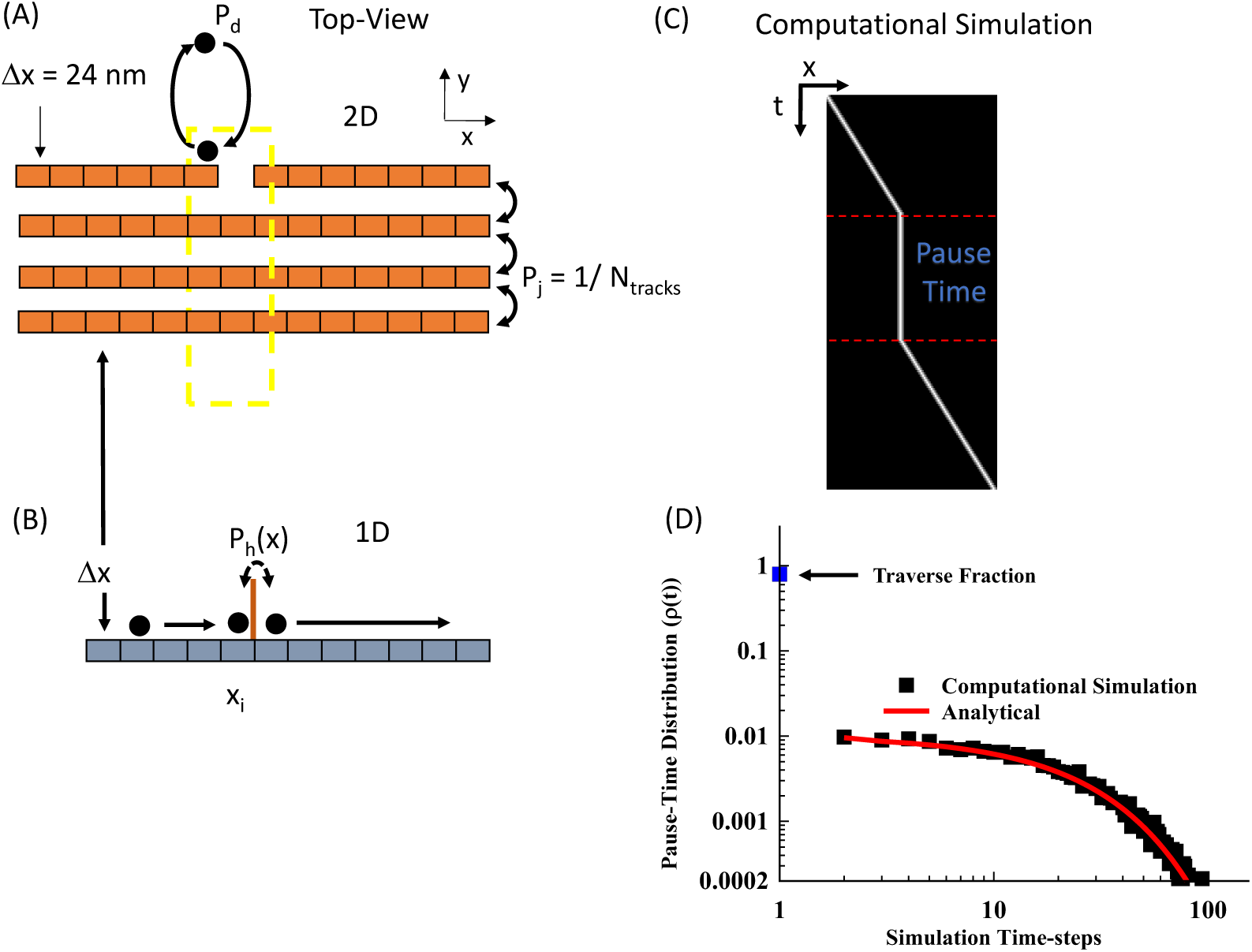
Dominant Motor-driven model. (A) We model an axonal bundle as lattice sites along a series of parallel tracks. Motors are modeled with a single detachment/attachment parameter (Pd). The probability a motor will jump between tracks is modeled as a uniform probability (Pj). MT-ends are modeled as missing lattice sites. (B) We reproduce kymograph results by making a one-dimensional simplification of our two-dimensional model. Each one-dimensional lattice site has the same spatial resolution as the two-dimensional lattice site (Δx). Vesicle inhibition by a MT-end site is now modeled as a probability (Ph), due to the loss of information about which track the vesicle is on. (C) A kymograph representation of a simulated vesicle pausing at the an x-lattice site, but with the exact track location is lost. (D) We introduce the Pause-time parameter to quantify and measure the probability a vesicle will pause at a MT-end. We show that our detachment parameter hypothesis (Red-line, equation A1) will show a uniform decrease in pause-time with increasing time. We compare our analytical expression with a computational simulation (Black-squares, 1000 simulations).

#### 3.4.2. Vesicle Mechanics Modelling

We model vesicle mechanics as a set of discrete processes, similar to a method previously used for single motor motility ^11^. In this model vesicle mechanics are coarse grained to a single rate-constant parameter, we call *pause-duration* (P_d_). This parameter represents vesicle pausing as a discrete finite probability threshold that includes: (i) vesicle detachment time from the microtubule; (ii) vesicle diffusion to the same or another MT; (iii) vesicle re-attachment time to a MT (Pd, Fig. 4A). All three processes occur on time-scales less than experimentally observable, including vesicle diffusion between tracks ^34^, and thus can be coarse-grained into a single parameter without losing accurate representation of experimentally observed results, which we show below.

Vesicles that detach at an MT end can hop between microtubule tracks in the bundle and reattach on any microtubule within the bundle (Fig. 4A). The probability of hopping on any microtubule track is equal for all tracks in the bundle (Pj = 1/Ntracks, where Ntracks is the number of tracks in a bundle). This equal weighting is assumed for simplicity in our model because experiments cannot distinguish any microtubule structure, but is also partially based on the experimental observation that axonal bundles are organized in a circular structure for smaller caliber axons ^31^. If a motor detaches it thus can re-attach to any nearest-neighbor on either side with symmetric equal weighting.

We follow a dynamic Monte Carlo simulation method to determine the vesicle behavior at each simulation time step (see Appendix A.3 for the algorithm). We define our minimum time-resolution window for simulations at 20 msec. The 20 msec limit is below the lower limit time-resolution of fluorescence microscopy experiments ^7,13^, but larger than single-motor stepping dynamics ^35^, which allows us to include ATP/ADP dynamics as a single parameter. At the beginning of each simulation, all probability parameters (Pd, Pj) are defined and fixed for the remainder of the simulation. At the beginning of each time step, random numbers are generated from an unweighted distribution. The vesicle is then determined to detach, hop, or walk by comparing the random numbers to their corresponding probability values.

#### 3.4.3. Modelling the Kymograph Representation

The vesicle mobility experiments presented above (Fig. 1) utilize a kymograph representation of experimental fluorescence microscopy images. The kymograph representation transposes two-dimensional intensity data onto a single one-dimensional line. Kymographs provide the ability to observe low-intensity events due to experimental imaging limitations in live-cells. However, as a consequence, kymographs cannot distinguish individual tracks within a bundle. We compare our model to experimental results by making an equivalent one-dimensional reduction of our two-dimensional simulation.

We reduce our two-dimensional model by representing each aligned lattice site on a track as a single one-dimensional lattice site (Fig. 4B). All two-dimensional lattice sites along individual tracks are aligned along the defined y-direction (Fig. 4A). If a motor is on any track in the bundle at a specific lattice site, then it is represented as along a single equivalent one-dimensional lattice site. This simplification preserves the original two-dimensional simulation results and spatial-resolution, while following the experimental kymograph method of representation. We also introduce a new parameter to account for bundle information lost due to experimental resolution limits, which we define as inhibition probability (P_h_) to model the probability a cargo will pause at a MT-end (Vertical line in Fig. 4B). The inhibition probability is mathematically defined as the number of microtubule ends divided by the number of microtubule tracks within the bundle at a lattice site (x). An example simulation shows an equivalent kymograph position as a function of time (Fig. 4C). The location of the motor along the x-axis of the bundle track (horizontal-axis) as a function of time also shows a pause at a MT-end (straight vertical line in Fig. 4C). The specific track the motor traverses is lost in the kymograph representation but pausing behavior at MT-ends is preserved.

#### 3.4.4. Pause-time Distributions Distinguish Vesicle Mechanics

To compare our model to experimental observations we focus exclusively on x-axis positions that include microtubule ends (the yellow dashed box in Fig. 4A). In this approach, the MT-end x-axis positions are identified and the time any simulated motor spends at that position is extracted. All times of all motors are binned in a histogram and normalized to obtain an ensemble distribution.

To accurately quantify and compare our model to experiment, we present a parameter that we call pause time (PT). The PT parameter measures the probability an ***aggregate distribution*** of motors will pause near a defect site in a bundle, similar to a previously developed observable used to describe single kinesin-1 motor motility on single microtubules with obstructions ^11^. The PT parameter does not measure the time on a specific microtubule, but quantifies how long a motor pauses at the same x-axis position as a microtubule end within the bundle (Fig. 4B), consistent with experimental limitations ^7^.

The advantage of the PT--distribution metric is that it can distinguish if a single rate constant (Pd) is sufficient to model how vesicles navigate MT-ends (see Appendix A.2 for derivations). First, vesicles that do not pause at a MT-end are only observed for one time-step/frame. Second, vesicles that pause have a probability to pause based on the pause-duration and inhibition probability. Both are represented mathematically as:

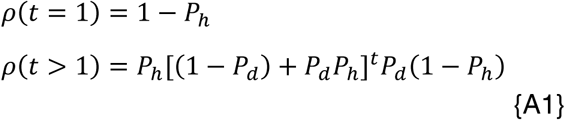

where *t* is in the units of the time-step/exposure interval and the probability of any specific vesicle pausing at an MT-end is P_h_≈1/N_track_, which assumes all tracks are equally accessible in the bundle. The number of time steps paused at the MT-end is denoted by t. This equation is also derived from a simple probabilistic model described in Appendix A.3.

An example distribution of simulated motors paused at a MT-end shows how to quantitatively determine a single rate constant model (Black squares in Fig. 4D). The probability that a motor/cargo will pause (y-axis) decreases with increasing pause-time (x-axis) for a simulated distribution. The single rate-constant probability model (eqn. A1) reproduces the distribution and provides specific testable parameters. We will use *both* eqn. A1 and computational models to distinguish how our model can explain the experimental results.

Lastly, to highlight the time-scale for a single-motor pause-times at MT-ends we modeled a pause-duration probability of P_d_=0.03 per 20 msec time-step, equivalent to an association rate-constant ∼1.5 sec^-1^, neglecting the time required for diffusion and attachment. For comparison, the canonical kinesin-1 motor on a bundled lattice has an dissociation rate-constant of ∼0.2 sec^-1^ (0.2 = 0.4 μm/sec / 2 μm, See discussion section for a table of comparisons) ^36^, while mammalian dynein/dynactin has an association rate-constant of ∼2 sec^−1 37^. We then calculated the PT-distribution for these motors at MT-ends (Black squares Fig. 4D).

### 3.5. Pause-time distributions for retrograde motion

Retrograde driven transport pausing at MT-ends exhibits a decreasing probability with exposure time. We used the same locations as measured for vesicle traverse fractions (Fig. 2), with the traverse fraction plotted as the intercept of the pause-time distribution (y-intercept Fig. 5). We aggregated the remaining pauses of the 359 events as a single distribution (Blue Squares, see Appendix 1 for method). The total probability distribution that retrograde driven vesicles will pause at any given MT-end was summed to be (0.24 +/− 0.02, or 1 – Traverse-fraction). The probability of pausing for a specific amount of time decreased from 0.06 +/− 0.005 for 110 msec to 0.0027 +/− 0.0027 for any time longer than 1 second (where the longer-time values is limited by the total number of events 1/359 = 0.0027). This decreasing pause-time distribution suggests that a single rate-constant determines the time any given vesicle will pause at a given MT-end. We note that multiple rate-processes may exist, but a two-rate model does not improve the qualitative or quantitative results compared to a single rate model (See Appendix A7).

**Figure 5:**
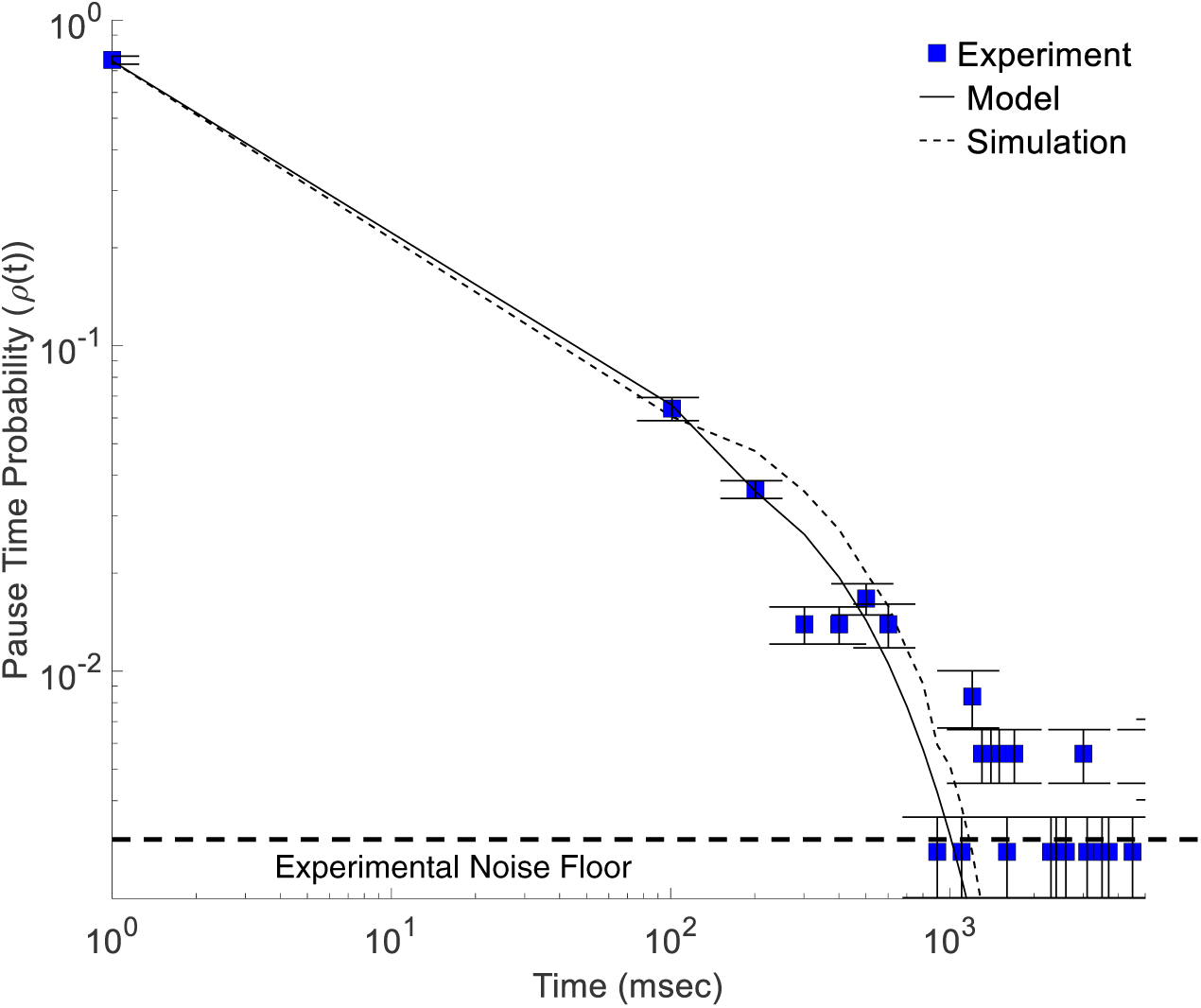
Retrograde-driven transport pause distribution as a function of time. Experimental data shows a decrease in pause-time probability with increasing time (Blue squares, N = 359). Both the analytical model (eqn. A1, solid line, Pd = 0.35) and the coarse-grained computational simulation (dashed line, Pd = 0.35) reproduce the observed experimental results.

We modeled the retrograde probability distribution based on our vesicle single pause-duration model (See Section 3.43 and Appendix A2). We set the vesicle traverse fraction in our model to the closest number integer MTs (N = 4) that fit the observed fraction in Fig. 2. We then determined the best parameter fit of our model to the data using un-weighted chi-squared analysis (See Appendix A6). Our model shows that the vesicle pause distribution follows a pause-duration probability of 0.35 per 110 msec (solid black line in Fig. 5).

It is important to compare our single-rate result with known protein dynamics involved with retrograde motion in order to understand the scale over which MT-end pausing mechanics occur. Retrograde motion is known to be driven by the molecular motor complex dynein/dynactin ^1,38^. We can compare our pause-duration to dynein/dynactin mechanics by comparing the association-time, which is the time a motor/vesicle is associated to the MT before dissociation (where dissociation-rate = 1 / association-time). Our model pause-duration probability is equivalent to a dissociation-rate of 3.9 sec^-1^ (k = -ln(1-.35)/0.11 sec), or an equivalent association time of ∼0.25 sec. Dynein-dynactin has a large variation in association times across species ^37^, with an association time between 32 sec for yeast ^38^, down to ∼2 sec for mammalian ^39^. Comparing association-times directly shows that the pause-duration at MT-ends is a factor of 4 or more shorter than for Dynein/dynactin association time. This suggests that Dynein/dynactin mechanics alone do not dictate how long vesicles will pause at MT-ends (See Discussion Section).

We computationally simulated vesicle mobility on a lattice with a single pause-duration probability parameter per time-step (see section 3.4, and Appendix 4 for model details). We reproduced experimental results by first modeling smaller time/space resolutions (24 nm spatial and 20 msec time resolution), followed by coarse-graining simulated results similar to the way camera exposure frames average. This coarse-graining approach results in the same general behavior, but increased variance (ref, Appendix A3, A5). We simulated vesicles with a Pd = 0.07 per 20 msec time-steps, which is equivalent to the experimental Pd = 0.35 for 100 msec exposure-time (where 0.35 = 0.07*100msec/20msec). This simulated pause-duration reproduced the experimentally observed pause-time distribution (dashed line). The difference between analytical and computational simulations are due to the slight differences caused by coarse-graining simulations (Appendix A5).

Both the analytical and computational models show that a single rate constant model accurately reproduces the majority of observed vesicle pause-times at MT-ends. We attempted to fit the longer pause-time pauses (>1 sec) with a model that assumes two possible distributions of pausing vesicles (See Appendix A7 for model); however, this two-rate model did not significantly improve the quality of fit, or the quantitative value of fit (Appendix A7). This is most likely because not enough vesicles were measured to accurately determine longer pause-times. Thus, a single constant Pd = 0.35 captures that majority of the observed pause-time distribution. This result thus suggests that a single pause-rate time-scale determines how long vesicles will pause at MT-ends during transport.

### 3.6. Pause-time distributions for anterograde motion

It is well known that retrograde and anterograde motion utilize fundamentally different motors to drive transport: with retrograde motion driven by dynein, and anterograde motion driven by kinesin. Dynein and kinesin family motors utilize different walking mechanisms ^40^. Further, different end-binding proteins localize at the minus or plus end of MTs ^41^. Consequently, it may be possible that retrograde and anterograde motion follow different transport in three following ways:

i. the difference in motor association/dissociate rates alone result in different mechanics at MT-ends. The difference in mechanics would result in fundamentally different observed pause-time distribution results.
ii. all the same motor/protein complexes co-bind on a vesicle and the MT to help navigate MT-ends, *but not result in a reversal*, which would result in a single pause-time distribution *regardless* of direction of travel.
iii. different proteins at plus/minus MT-ends may mediate vesicle navigation of MT-ends and dictate vesicle pause-duration, regardless of the motor. These different proteins would then result in different pause-durations for anterograde and retrograde motion. Further, this possibility would be distinguished from (i) if vesicle pause-durations do not match individual motor dissociation/association rate-constants.

We sought to test these three hypotheses by quantifying the pause-time distribution for anterograde motion separate from retrograde motion.

Anterograde driven transport pausing at MT-ends exhibits a continuously decreasing probability with time. We calculated the distribution of Anterograde pausing at MT-ends by aggregating pauses of 388 events from 15 different MT-ends in 4 different experiments (see Appendix A1). The probability of pausing for a specific amount of time decreased from 0.1 +/− 0.005 for 110 msec to 0.0025 +/− 0.0025 for any time longer than 1 second (where the longer-time values is limited by the total number of events 1/388 = 0.0025).

The pause-time distribution follows the same general reduction as a function of time observed for retrograde driven transport. This suggests that our overall single rate-constant model can describe both directions of motion. Further, anterograde and retrograde driven pause-time distributions are significantly different from each other. However, we must still model anterograde pause-times in order to distinguish between hypotheses (i) and (iii).

We modeled the anterograde probability distribution based on the same attachment/detachment mechanics (See Appendix A2). We set the vesicle traverse fraction in our model to the observed fraction in Fig. 2. We set the vesicle traverse fraction in our model to the closest number integer MTs (N = 4) that fit the observed fraction in Fig. 2. We then determined the best parameter fit of our model to the data using un-weighted chi-squared analysis (See Appendix A6). Our model shows that the vesicle pause-time distribution follows a pause-duration of 0.5 per 110 msec (solid black line in Fig. 6). Biasing our model toward earlier pause-times or later pause times did not significantly improve the fit to data (Appendix A6). Thus, a single constant P_d_ = 0.5 +/− 0.1 captures that majority of the observed pause-time distribution.

**Figure 6:**
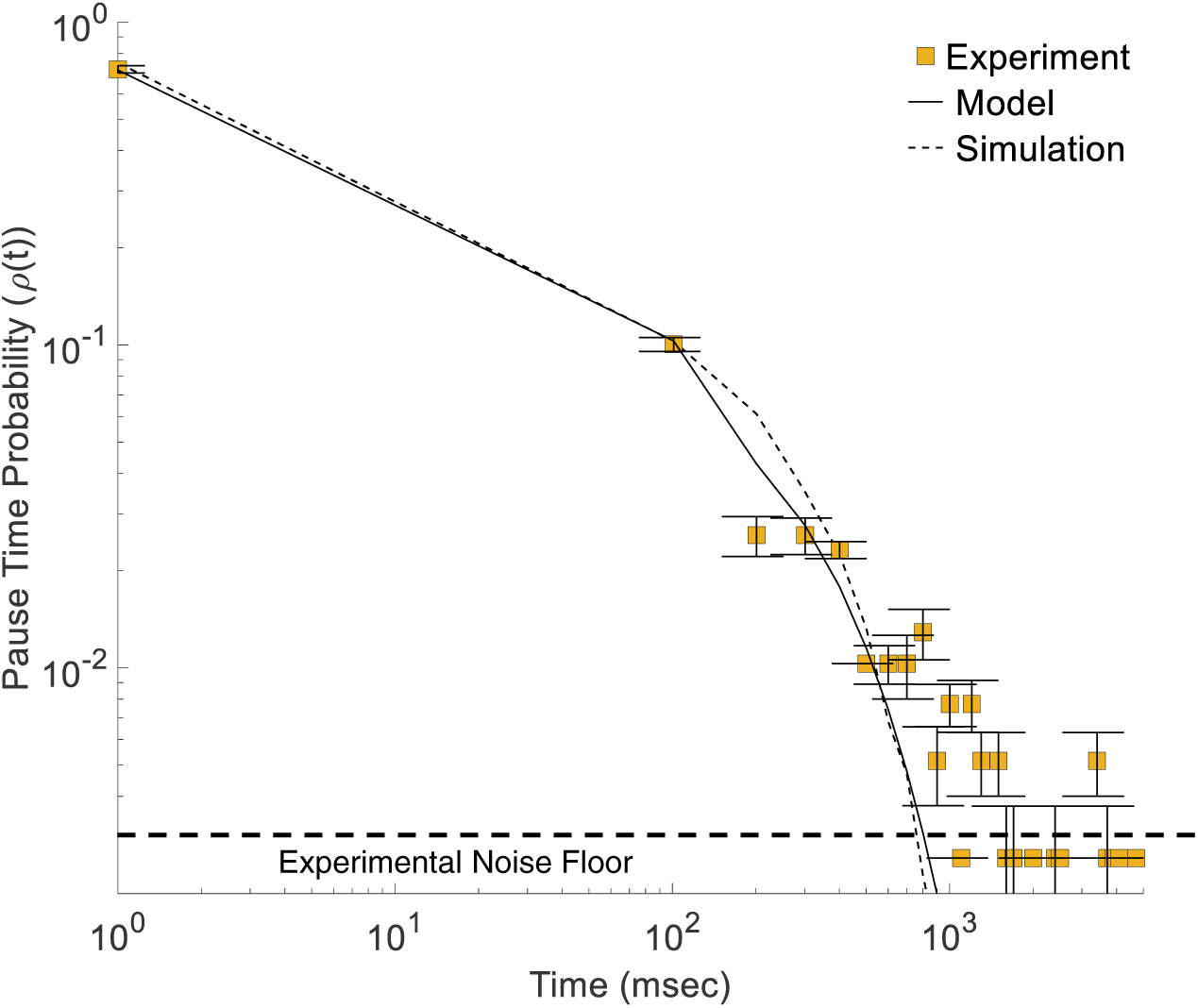
anterograde-driven transport pause distribution as a function of time. Experimental data shows a decrease in pause-time probability with increasing time (Yellow squares, N = 388). Both the analytical model (eqn. A1, solid line, Pd = 0.5) and the coarse-grained computational simulation (dashed line, Pd = 0.65) reproduce the observed experimental results.

To distinguish between hypotheses (i) and (iii), it is important to compare our observed pause-duration results to the known anterograde motor protein in order to understand the time-scale of vesicle pausing, as done above with Dynein/dynactin. Our anterograde pause-duration probability is equivalent to an off-rate of 6.3 sec^-1^ (k = -ln(1-.5)/0.11 sec), or association-time of ∼0.15 sec. UNC-104 is the known kinesin motor the drives anterograde synaptic vesicle precursor ^42^ motion. UNC-104 has a known association-time of ∼1-2 sec ^43^, which is equivalent to an off-rate of 0.5 – 1 sec^-1^. Thus, our observed pause-duration time-scale at MT-ends is a factor of 3-6 times faster than known single motor UNC-104 detachment kinetics. This comparison suggests that single motor mechanics alone does not determine vesicle pause-times at MT-ends, ruling out hypothesis (i) above (See Discussion Section).

We computationally simulated vesicle mobility for anterograde motion exactly the same as with retrograde motion (discussed above) but with a different pause-duration probability (see section 3.4, and Appendix A2 - A4 for model details). Our simulated vesicles with a P_d_ = 0.13 (equivalent to an experimental P_d_ = 0.65 in 110 msec coarse-grained time) reproduced experimentally observed pause-time distribution (dashed line). The difference between analytical and computational simulations are due to the slight differences caused by coarse-graining simulations.

The difference between anterograde and retrograde pause-time distributions, combined with the difference in time-scales between pause-duration and single motor association/dissociation times, suggests that vesicle pause-times are determined by different proteins interacting with the vesicle at the time it reaches a MT-end (hypothesis (iii)). Further, retrograde driven motion has a lower pause-duration probability than anterograde driven motion (Pd = 0.35 for retrograde, and Pd = 0.5 for anterograde). This difference is consistent with previous observations of pausing at MT-ends^22,21,20,21,14^, *and* compares with the difference in single motor mechanics experiments that show dynein motors have a longer association time >2 sec ^39^, than < 2 sec for UNC-104 ^43^.

## 4. Conclusions

We have quantified the pausing behavior of synaptic vesicle precursors at microtubule-end locations (MT-end) in vivo. We showed that vesicles have the same probability of being stopped at MT-end locations for both anterograde and retrograde motion. We also showed that vesicle reversals are a small fraction of motion after vesicles leave MT-end locations, suggesting that tug-of-war mechanics are a minority of behavior. We propose a single-rate model by which vesicles navigate MT-ends, based on the observation that vesicles pause for different times depending on direction of motion before reaching a MT-end. We show that this model reproduces both anterograde and retrograde pause-time distributions. We also computationally simulate vesicles at MT-ends with a single-rate pause-duration parameter and show that the simulations reproduce observed experimental pause-times.

## 5. Discussion

### 5.1. Implications of a Single Rate-Constant analysis and model

The results in this study show that the direction of travel at the time the vesicle reaches a MT-end dominates how long the vesicle pauses before continuing its motion. Combined with the low reversal fraction at MT-ends, this direction dependent pausing suggests that vesicles rely on mechanics of the driving motor in combination with other possible co-binding proteins to navigate MT-ends. Our pause-time analysis and single rate-constant result is significant because different mechanisms can occur simultaneously affecting the qualitative nature of the pause-time distribution. For example a UNC-104 motor mutation may result in sub-populations of longer vesicle pausing at MT-ends, which in our model would be described by more than one rate-constant, but not result in a change in the average pause-time; any interpretation based on average pause-time alone would miss the difference in mechanics observed by measuring pause-time distributions. Thus, measuring the pause-time distribution and comparing to a rate-constant model provides increased understanding of the underling mechanics.

### 5.2. Comparison of Pause-duration at MT-ends to Single Motor Mechanics

A direct comparison between our observed pause-durations and single motor mechanics can provide insight into the possible contribution motor mechanics may have in vesicle navigation of MT-ends. Pause-duration time scales for both anterograde and retrograde motion are also significantly shorter than would be expected for single motor motility. We directly compare our observed pause-duration rates to known single motor detachment rates (Table 1). Single molecular motors engage in motility for a finite time before detaching from a filament. This association time can be converted to a *detachment rate* by dividing the velocity by the run-length, both of which are easily quantified by single molecule experiments. This detachment rate is then directly comparable to our pause-duration because both quantify the time until an event occurs.

**Table 1:**
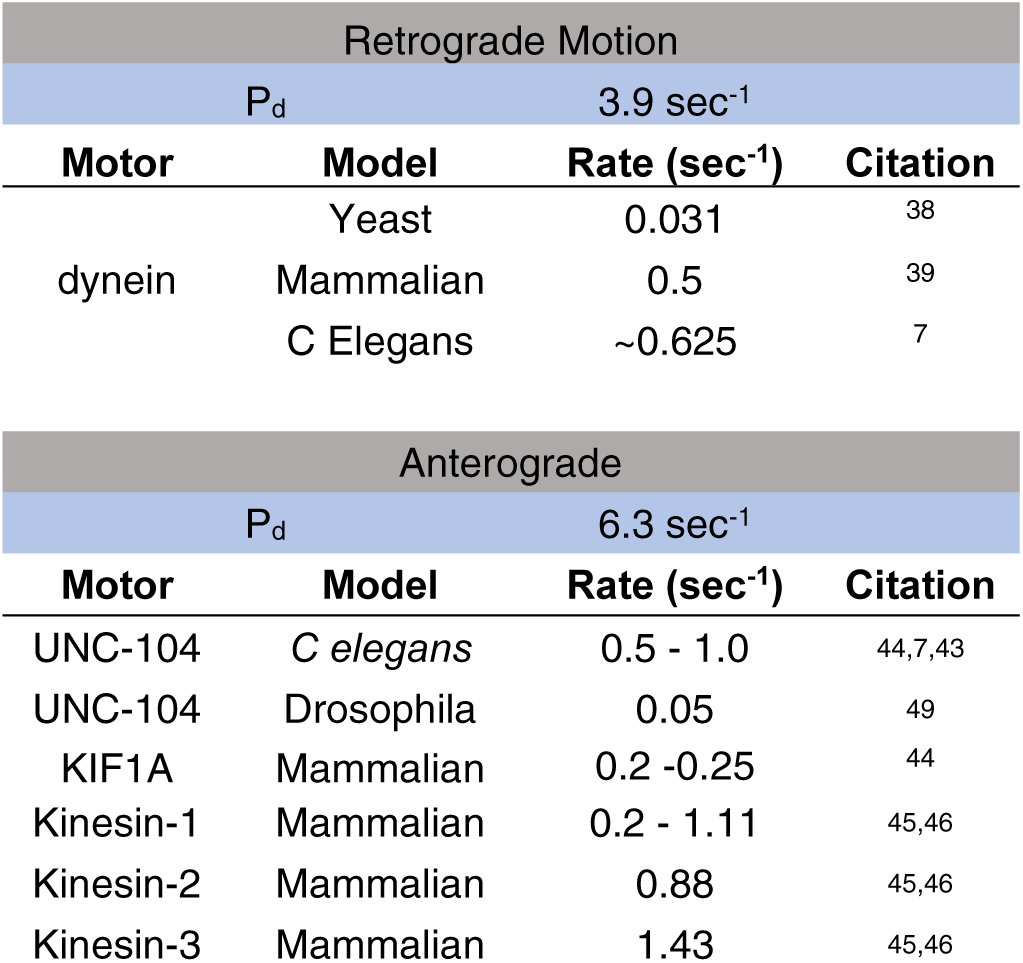
Comparison of Pause Duration and Single motor detachment rates for known SVP motors

| Retrograde Motion |  |  |  |
| --- | --- | --- | --- |
| P<sub>d</sub> |  | 3.9 sec<sup>-1</sup> |  |
| Motor | Model | Rate (sec<sup>-1</sup>) | Citation |
| dynein | Yeast | 0.031 | 38 |
|  | Mammalian | 0.5 | 39 |
|  | C Elegans | ~0.625 | 7 |
| Anterograde |  |  |  |
| P<sub>d</sub> |  | 6.3 sec<sup>-1</sup> |  |
| Motor | Model | Rate (sec<sup>-1</sup>) | Citation |
| UNC-104 | *C elegans* | 0.5 - 1.0 | 44,7,43 |
| UNC-104 | Drosophila | 0.05 | 49 |
| KIF1A | Mammalian | 0.2 -0.25 | 44 |
| Kinesin-1 | Mammalian | 0.2 - 1.11 | 45,46 |
| Kinesin-2 | Mammalian | 0.88 | 45,46 |
| Kinesin-3 | Mammalian | 1.43 | 45,46 |

Both retrograde and anterograde pause durations are orders of magnitude faster than single motor detachment rates. Dynein, which drives retrograde motion, exhibits a factor of 5-8 times slower detachment-rates for mammalian and *C elegans* models than our observed pause-duration ^7^. Kinesins, which drive anterograde motion, all exhibit a 5-10 times slower detachment rate than our observed pause-duration ^44 45,46^. The time-scale difference between single motor detachment-rates and vesicle pause-durations strongly suggest that single motor mechanics contribute a subordinated component to how vesicles navigate MT-ends.

### 5.3. Multi-protein interactions at MT-ends

Our current results show that a single rate-constant model describes vesicle pause behavior at MT-ends but does not distinguish if more than one protein is involved in the process. It is well established that multiple protein process pathways can work in parallel resulting in a single observed rate-constant ^47^. While the difference in average rate can distinguish that different mechanics exist for retrograde and anterograde motion, but average rate-constant result *alone* cannot distinguish if multiple proteins coordinate at MT-ends.

*In combination with future mutation studies*, our pause-time distribution metric and single-rate model can be used to distinguish the existence and mechanism of multi-protein interactions at MT-ends. If another protein is involved with the process by which vesicles navigate MT-ends then altering or removing that protein should significantly alter the average pause-time distribution. Specifically, if a protein enhances the ability for vesicles to navigate MT-ends, then removing that protein would result in a smaller detachment probability, and equivalently a longer pause-time. Conversely, if a protein reduces the ability of a vesicle to navigate a MT-end then removing the protein would increase the detachment probability, or equivalently result in a shorter pause-time. Future experiments could measure pause-time distributions under conditions with known proteins that alter or modulate MTs.

### 5.4. Role of MT-ends in long-time/distance processes

Many cell signaling processes rely on transport of signals from the periphery of the axon to the cell nucleus, or vice versa with a signal sent from the cell nucleus to the periphery. Vesicles must navigate any MT-end they encounter during that process. While each pause at a MT-end does not contribute a significant amount to the overall transport time, accumulating multiple vesicle pauses along the axon can contribute significantly to transport times. Further, if a signaling process requires tightly controlled transport times then any deviation can result in altered cellular behavior. *The results in this study show that the mechanics of the driving motor are important in distinguishing the amount of time navigating MT-ends contributes to overall transport time*.

### 5.5. Difference between Pause-times in long-time/distance processes

The difference in pause-time distributions at MT-ends for retrograde and anterograde motion has significant implications for long-range axonal transport. Retrograde driven pause-times showed a significantly longer pause-time than for anterograde motion. This difference suggests that each vesicle pause at a MT-end during retrograde motion will contribute more to axonal transport than anterograde motion. If vesicle transport times are increased by MT-ends, *the results in this study suggest that retrograde and anterograde axonal transport times will be differentially altered by MT-ends*. Future studies could begin to explore the contribution of differential MT-end pausing on long-range cell signaling and disease.

## 6. Acknowledgements

## 8. Appendix

### Appendix A1: Data Aggregation and error methods

We quantified aggregate pause-time distribution for MT-ends by combining all motor pauses into a single binned distribution using the following method (presented as an example in Table CC below):

i. We binned cargo based on the number of frames they were observed at the x-position along the kymograph such that: cargo that traversed a MT-end were observed for 1-frame, cargo that paused for 1-exposure-time were observed for 2-frames, etc …
ii. We distinguished anterograde and retrograde cargo motion and counted them separate bins.
iii. We performed [i] and [ii] for 15 different MT-end locations across 4 different experiments.
iv. We then summed the individual bins into a single final bin as follows: all cargo that traverse MT-end locations (defined as bin = 1-frame discussed in [i]) were summed together for a total of 288 tracks for anterograde and 271 tracks for retrograde; all cargo that paused for a single exposure-time (defined as bin = 2-frames discussed in [i]) were summed together for a total of 39 tracks anterograde and 23 tracks for retrograde; etc..
v. The number of vesicles in each bin are then normalized by the total number of tracks 359 tracks anterograde and 388 tracks for retrograde;
vi. The resulting distributions are plotted in Fig. 5 for retrograde and Fig. 6 for anterograde
vii. Errors were calculated as the standard-deviation in number of vesicles across all MT-ends, for a given bin (n = 1, 2, 3, …) divided by the total number of vesicles observed.

**Table A1:**
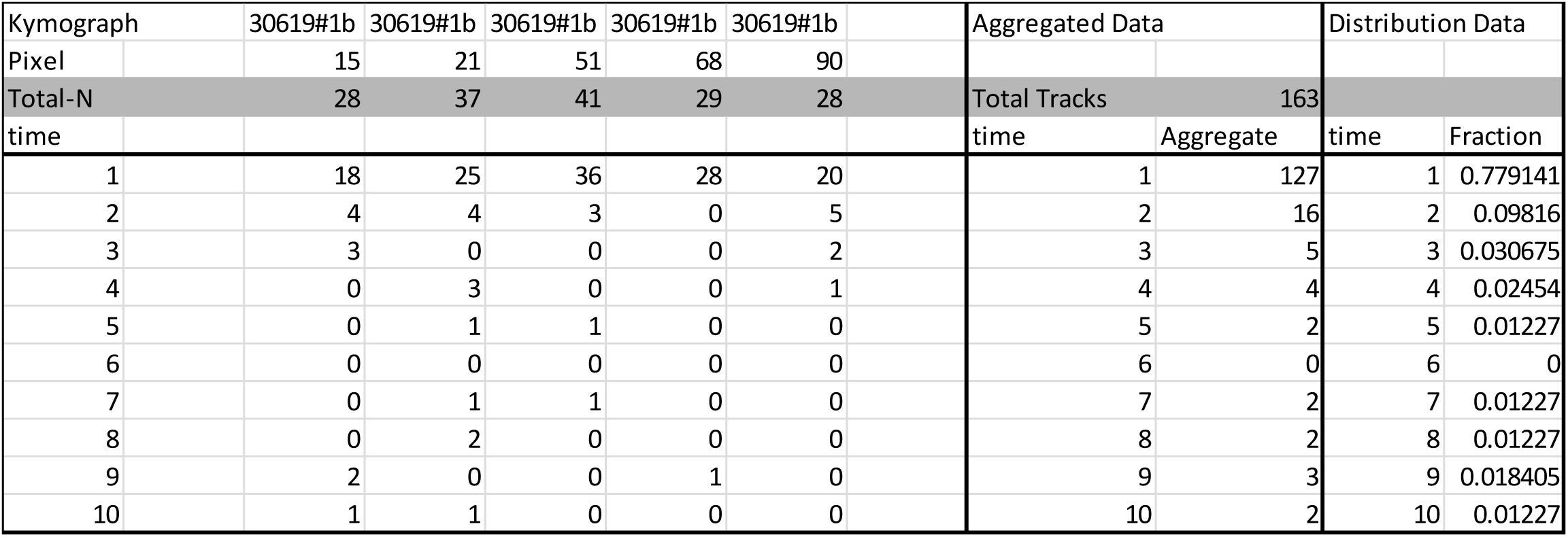
Anterograde pause distribution Aggregation Method Example

**Figure A1:**
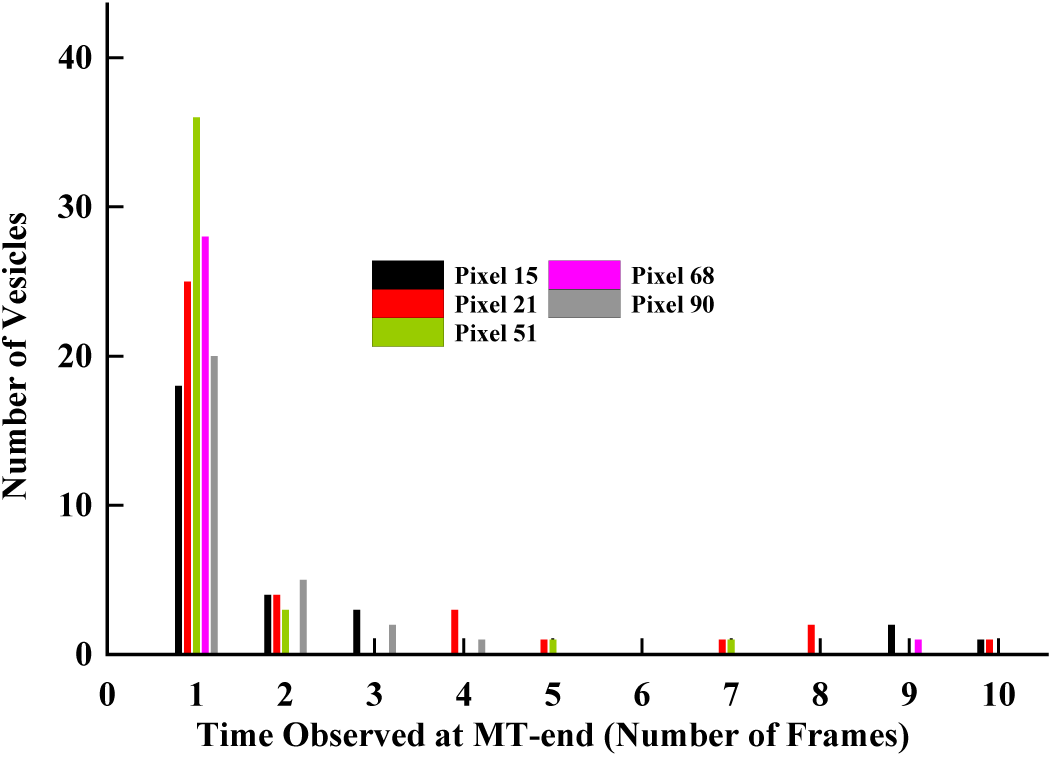
Anterograde pause distribution Aggregation Histogram.

### Appendix A2: Derivation of Probabilistic Model of Motor Pause Distributions at Microtubule ends

**Figure A3:**
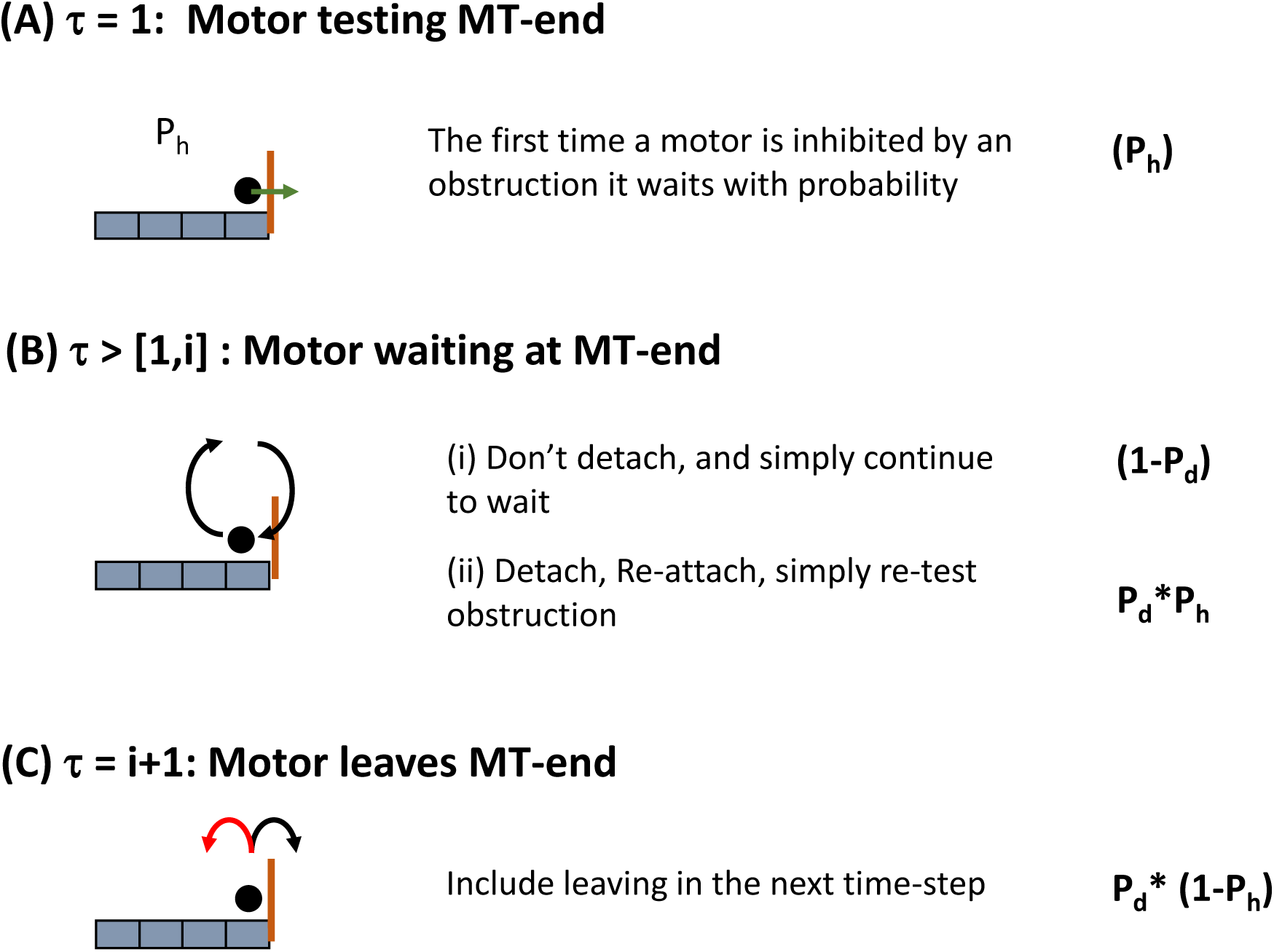
Analytical Probability Model. (A) The probability a motor pauses at a site with an obstruction (Ph) is equal to the number of ends divided by the number of tracks. (B) The probability a motor waits a single time step at an obstruction is the combined probability of not detaching and the probability that a motor detaches and re-attaches to the same track. (C) The probability a motor leaves in a time step is the probability that a motor detaches, re-attaches, and occupies a new track.

We now derive the analytical model of single motor pausing at microtubule ends *in the one-dimensional simplification*. This derivation works from the experimental perspective that we do not know the individual microtubule track a motor is on. Further, the derivation is based on a combinatorial approach to a one-dimensional experimental observation. At each time-step a motor may execute a combination of different processes each one dependent upon the other in the following ways:

i. The first time a motor reaches a microtubule end position there is probability it may ***not*** be on the same microtubule track as the microtubule end (1 – P_h_), and thus ***not pause***.
ii. If the motor is on a microtubule track with a microtubule end then it will pause for its first time-step at that position (P_h_), as shown in Fig. A.2A.
iii. If a motor has paused from condition (ii) then there are two ways it may continue to pause (Fig. A.2B):
  a. The motor doesn’t detach (1 – P_d_)
  b. The motor does detach, re-attach, and is still on the same track as the microtubule end (P_d_ P_h_)
iv. The motor will continue testing condition (iii) above for (t) time-steps, with each step multiplied by the previous number of attempts
v. At (t) time-steps a motor detaches and re-attaches to a different microtubule track than the microtubule end (P_d_ (1 – P_h_)), as shown in Fig. A.2C.

The first process above (i) is simply the number of motors that do not pause at a known microtubule end site. The following processes (ii)-(iv) are the number of possible ways motors may test a microtubule end site and combine into eqn. 1. We have included the time required for a motor to re-attach to another microtubule track (iv) because this would be how experiments would determine if a motor has been able to traverse a microtubule end.

### Appendix A3: Vesicle Simulation Algorithm

**Figure A4:**
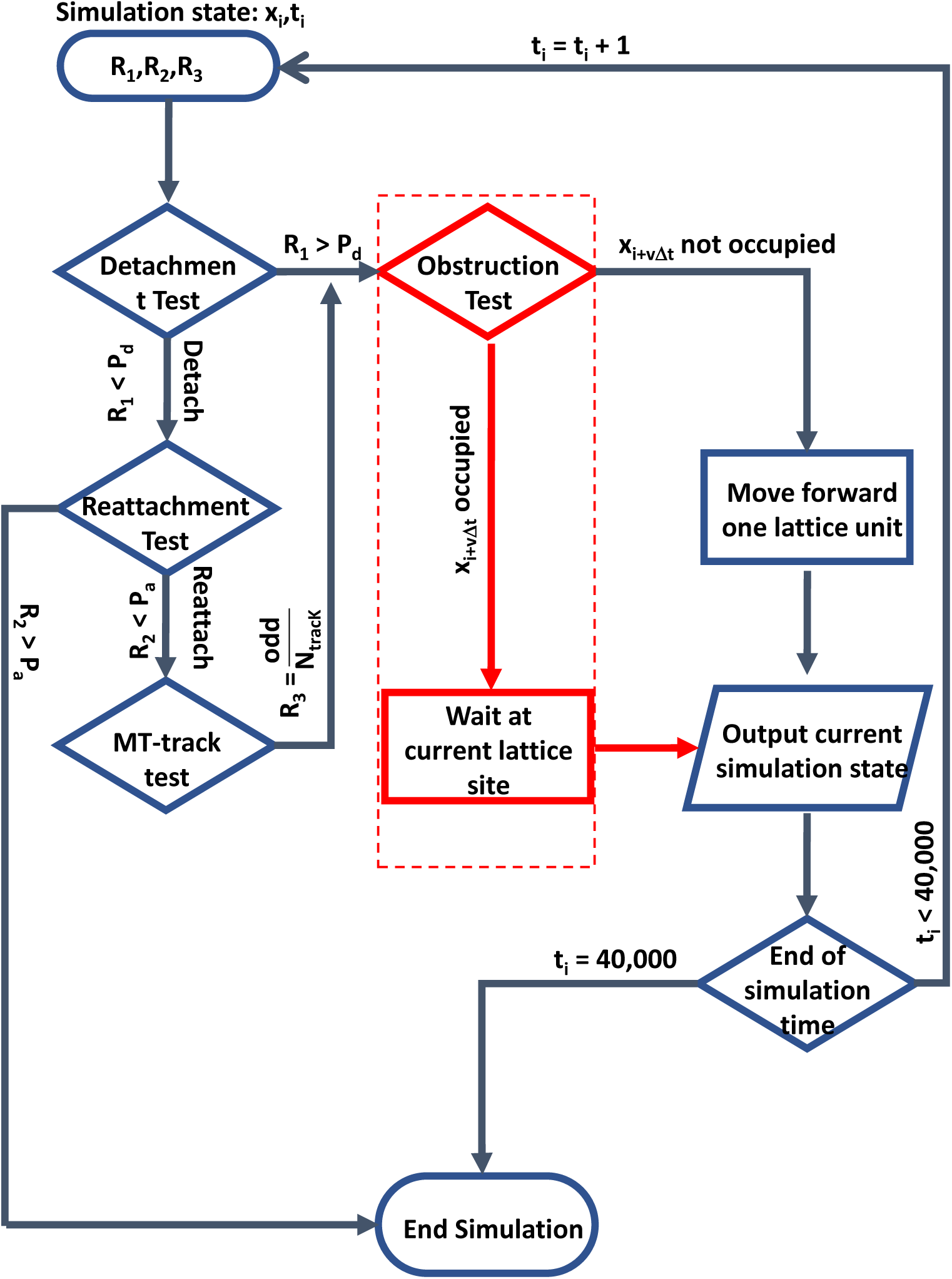
Simulated Vesicle Algorithm Flow Chart.

The vesicle decisions at each time step (ti) follow a specific algorithm, shown in Fig. A3. At the beginning of each simulation, a bundle generated from the bundle algorithm is loaded from a file into a two-dimensional array U. The motor simulation then uses the fixed obstruction locations for the remainder of the simulation.

First, three random numbers (R1, R2, R3) are chosen at the beginning of each time step, chosen from an unweighted python number generator random.rand(), which uses a Mersenne Twister algorithm to generate random numbers between [0, 1). Following the dynamic Monte Carlo approach, all random numbers are discarded at the end of each time step and a new set of random numbers is chosen at the next time step.

Second, vesicle detachment, re-attachment, and microtubule-switching tests were run, shown in Fig. A3. A motor detaches if R1<Pd, otherwise the obstruction test is run. If a vesicle detaches, then it re-attaches in the same time-step.

Third, a re-attached vesicle randomly chooses a microtubule track to hop on, determined by the random number R3, with all tracks equally likely to be chosen. The algorithm to determine a specific track is as follows: the probability for all tracks are assigned a decimal range (i.e. Track 1 P1= [0,1/Ntrack), Track 2 P2= [2/Ntrack,1/Ntrack)), a vesicle reattaches to a track if R3 is within the assigned range (i.e., if R3 ϵ P1, then the motor is attached to Track 1).

Once motor-bundle interactions have been determined, a motor-obstructions test is run, shown in Fig.A2. If there is no motor obstruction at the next lattice site to be occupied by the motor, then the motor moves to that site. If there is an obstruction at the next lattice site, then the motor remains on the same lattice site for the next time step.

The simulation ends if the motor reaches the end of the bundle or the time steps exceed a maximum threshold. In the case of perfectly aligned bundles, motors always reach the end of the bundle. In the case of completely unbiased bundles, motors may not reach the end of the bundle depending on the number of obstructions and their respective inhibition probability. Therefore, a maximum threshold of 10,000 time steps was set.

Notably, alternative algorithms, such as the Gillespie Algorithm, have been used to model single-motor motility. However, these algorithms would result in the equivalent measurable quantities to those described in this manuscript. Our interest is to provide an algorithm that mimics the randomness at the local time step level, which we can easily achieve with dynamic Monte Carlo simulation.

### Appendix A4: Coarse-Graining Method

Vesicles were coarse-grained (CG) after 2D bundle motility simulation and a 1D simplification. Simulations were CG by defining a new lattice site, which is an integer number of un-coarse-grained lattice sites (ΔX = n*Δx). The time in the CG lattice site is calculated as the difference in time a motor spends within the n-lattice sites, i.e., the CG simulation time for each ΔX lattice site. The CG inhibition probability (Ph) is the simple sum of all Ph values within the n-lattice sites, which is based on the assumption that each time a motor encounters a Ph, it has no memory of any previous microtubule end.

It is helpful to distinguish different conditions that lead to different CG simulation times at each CG lattice site to better understand how they may be observed experimentally.

For the completely biased bundle, there are two conditions:

i. A motor hops through all un-CG lattice sites without obstructions and the CG time is ΔX/Δx;
ii. A motor pauses at one or more un-CG lattice microtubule end sites within a single CG-lattice site, resulting in a CG-time of ΔX/Δx + Pause-Times at microtubule ends. Note that the Ph is the sum of all microtubule ends within a CG-lattice site and thus corresponds to the increased CG time.

We note that simulation time steps and CG time can be converted to the Lab-time. In the un-CG simulation, a single time step is 30 msec. CG time steps are integer multiples of un-CG time-steps (n*30 msec, n = 1, 2, 3, etc.). Thus, the Lab-times used in this simulation are integer multiples of 30 msec.

### Appendix A5: Pause Time Measurements Depend on Camera Exposure Times

**Figure A5:**
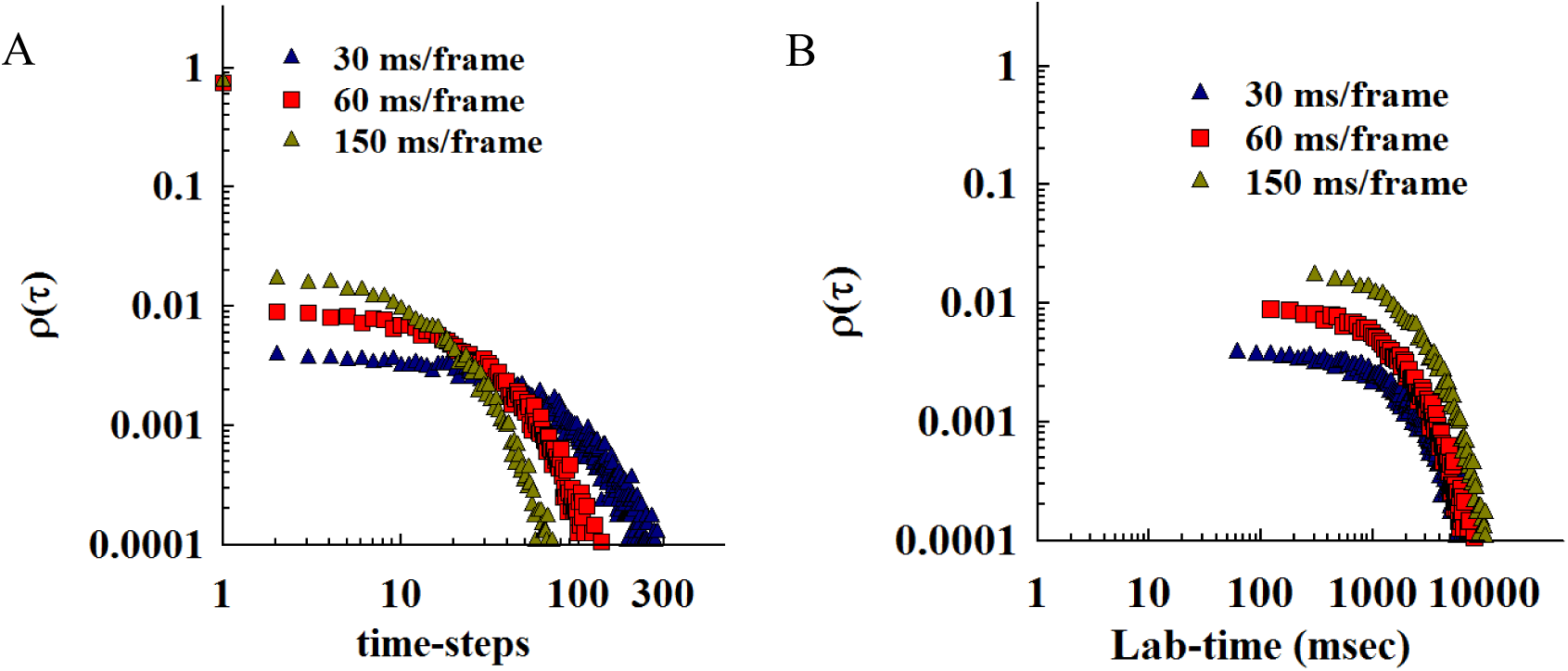
Pause-time distributions must be determined in Lab-time: (A) Pause-time distributions are calculated for 1000 simulations of vesicles with Pd = 0.03 per time-step equivalent to 30 msec/frame camera exposure (Blue triangles). The same simulated vesicles were coarse-grained by 2-frames (Red squares) and 5-frames (Green squares). Plotting the pause-time distributions on time-step scales appears to change the distributions. (B) The same pause-time distributions in (A) converted to lab-time. All distributions show the same qualitatively equivalent distributions, but with different density of points.

The resolution of the experimental camera exposure time affects the ability to measure PT. This phenomenon has been of considerable interest in many single-molecule experiments focusing on improving instrumentation ^48,49^ and data analysis techniques ^50^. More recently, there has been interest in de-convolving real rate-processes from the camera response at or near the single-frame resolution limit ^51^. Considering the interest in such resolution limitations, we show that Pause-time distributions must be measured in lab-time frames rather than experimental camera exposure time.

We use a single bundle simulation with vesicles that have the same detachment (P_d_ = 0.03). After the simulations, we calculated the one-dimensional bundle (See Section 3.4). We then coarse-grained the one-dimensional results by combining a number of x-lattice sites (1, 2, 5 bins, see appendix A.4 for algorithm). We then correlate the increased spatial limit with increase time resolution (1, 2, 5 time steps) to represent different camera exposure times (30, 60, and 150 msec). This method mixes time/space resolution limits, which is consistent with experimental experiments.

### Appendix A6: Chi-squared analysis of model versus data

**Table A6:**
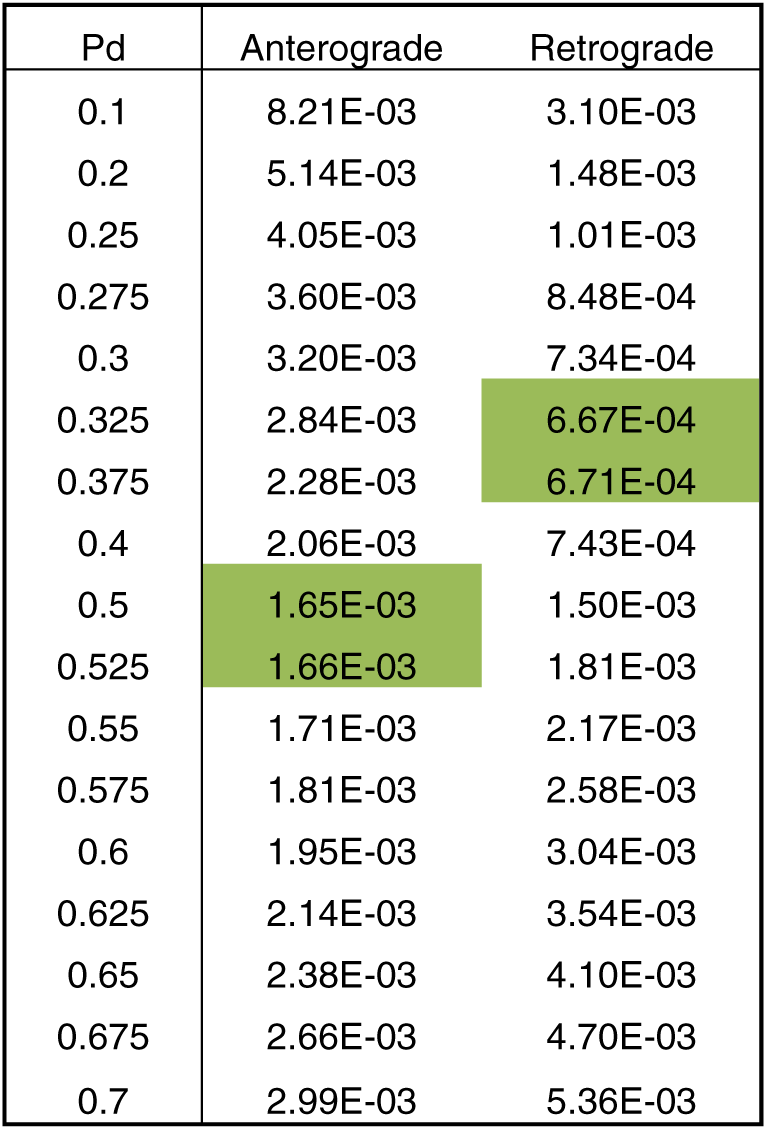
Chi-square values for model/data comparisons

### Appendix A7: Two-rate Model Fit to Data

We fit the observed anterograde pause-time distribution to a two-rate model to improve the quality of fit to longer pause-times. We observed that our single-rate model reproduced early time pauses but did not match the longer-time pauses (Fig. 5, 6). We hypothesized that the observed pause-time distribution may be a collection of more than one population of vesicle pause-durations. If this hypothesis is true, then the observed pause-time distribution may be best fit with a model that includes a mixture of two different pause-duration parameters.

We tested this hypothesis by making a two-rate model to fit to the anterograde pause-time distribution. We created our two-rate model as follows:

i. We made two separate models (ρ_1_, ρ_2_), each described by eqn. A1 in section 3.4.3, with a single fixed pause-duration (P_d1_, P_d2_). For example, we made two models with P_d1_ = 0.1 and P_d2_ = 0.7.
ii. We then added a variable parameter (ϕ) that determined the fraction that each model would contribute in the two-rate fit, but constrained to a combined fraction summed to 1.
iii. We then summed the fractional contribution from each model. This fractional sum was then compared to the observed pause-time distribution.
iv. Finally, we used a chi-square analysis to determine the best fraction value that would fit given the two pause-duration values.

This two-model fit is best described mathematically as:

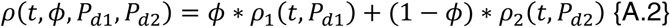

**Figure A8:**
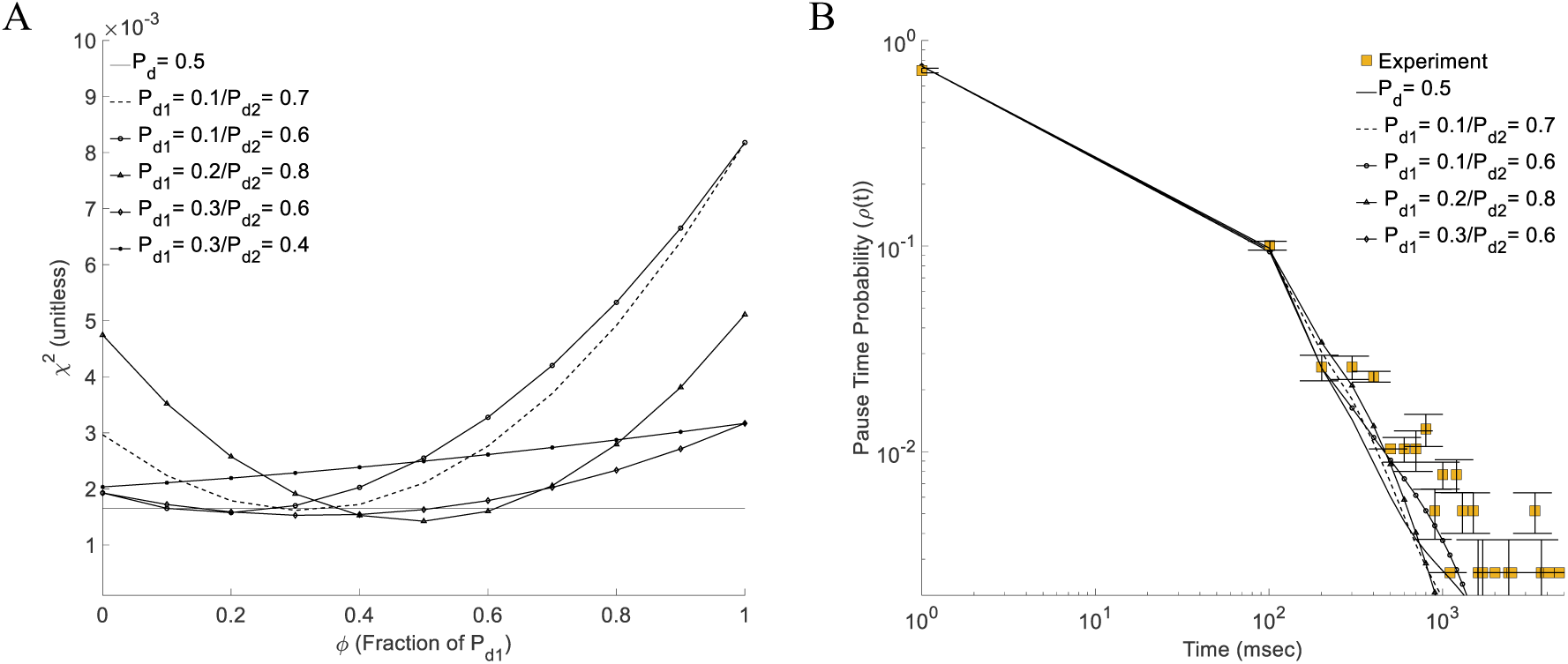
Anterograde pause distribution fit with a mixed model. (A) Chi-Squared analysis of different model fits to anterograde data. Mixed models were created as a fraction of two different pause-duration parameters (See text in Appendix A7), calculated based on the analytical model in Appendix A3. The chi-squared values were calculated as a function of the fraction parameter in the model (ϕ, x-axis). (B) The lowest chi-square fits for each mixed model are plotted as compared to experimental data (Same line/symbols used as in A). No mixed model significantly improves the quality of fit as compared to a single pause-duration parameter model.

We fit the two-rate model described above to observed anterograde pause-time distribution (Fig. A8). We created a combination of different pause-duration values with a low-value (P_d1_) and a high-value (P_d2_). We varied the fraction of each low/high value and compared the combined two-rate model to observed data using the same unweighted chi-squared analysis for the single-rate model. Finally, we plotted the different chi-squared fits as a function of the fraction-parameter (Fig. A8 A).

No two-rate model significantly improved the chi-squared fit as compared to the single-rate model fit (Fig. A8 A). The lowest chi-square value for each two-rate fit matched the single-rate fit (solid horizonal line). The best two-model fit (P_d1_ = 0.2/ P_d2_ = 0.8) gave ∼13% improved fit over the single-rate model. Further, the two-rate model (P_d1_ = 0.3/ P_d2_ = 0.4) gave the worst chi-squared fit, with the best chi-squared occurring when the model is entirely the single-rate of 0.4 (ϕ = 0); this is likely due to both pause-duration parameters below the 0.5 value for the single-rate model. *These results show that a two-rate model does not significantly improves the quantitative fit to observed anterograde pause-time distribution.*

The two-rate model slightly improved the qualitative fit as compared to the single-rate model (Fig. A8 B). We compared the best-fit pause-time distributions for different two-rate models, obtained using chi-square analysis above (Fig. A8 A), to the observed anterograde pause-time distribution (Squares Fig. A8 B). We also compared the two-rate models with the best-fit single-rate model (P_d_ = 0.5, Solid line Fig. A8 B). *All two-rate models slightly improved the quality of fit to longer-time pauses but at the expense of worse fits to short-time pauses*.

## Notes

### Competing Interest Statement

The authors have declared no competing interest.

