## Supplementary Table 2 for "Vesicle Navigation of Microtubule Ends Distinguished by A Single Rate-Constant Model"

| Pd | Anterograde | Retrograde |
|---|---|---|
| 0.1 | 8.21E-03 | 3.10E-03 |
| 0.2 | 5.14E-03 | 1.48E-03 |
| 0.25 | 4.05E-03 | 1.01E-03 |
| 0.275 | 3.60E-03 | 8.48E-04 |
| 0.3 | 3.20E-03 | 7.34E-04 |
| 0.325 | 2.84E-03 | 6.67E-04 |
| 0.375 | 2.28E-03 | 6.71E-04 |
| 0.4 | 2.06E-03 | 7.43E-04 |
| 0.5 | 1.65E-03 | 1.50E-03 |
| 0.525 | 1.66E-03 | 1.81E-03 |
| 0.55 | 1.71E-03 | 2.17E-03 |
| 0.575 | 1.81E-03 | 2.58E-03 |
| 0.6 | 1.95E-03 | 3.04E-03 |
| 0.625 | 2.14E-03 | 3.54E-03 |
| 0.65 | 2.38E-03 | 4.10E-03 |
| 0.675 | 2.66E-03 | 4.70E-03 |
| 0.7 | 2.99E-03 | 5.36E-03 |
