## Supplementary figures and images for "Vesicle Navigation of Microtubule Ends Distinguished by A Single Rate-Constant Model"

### Supplementary Figure 1

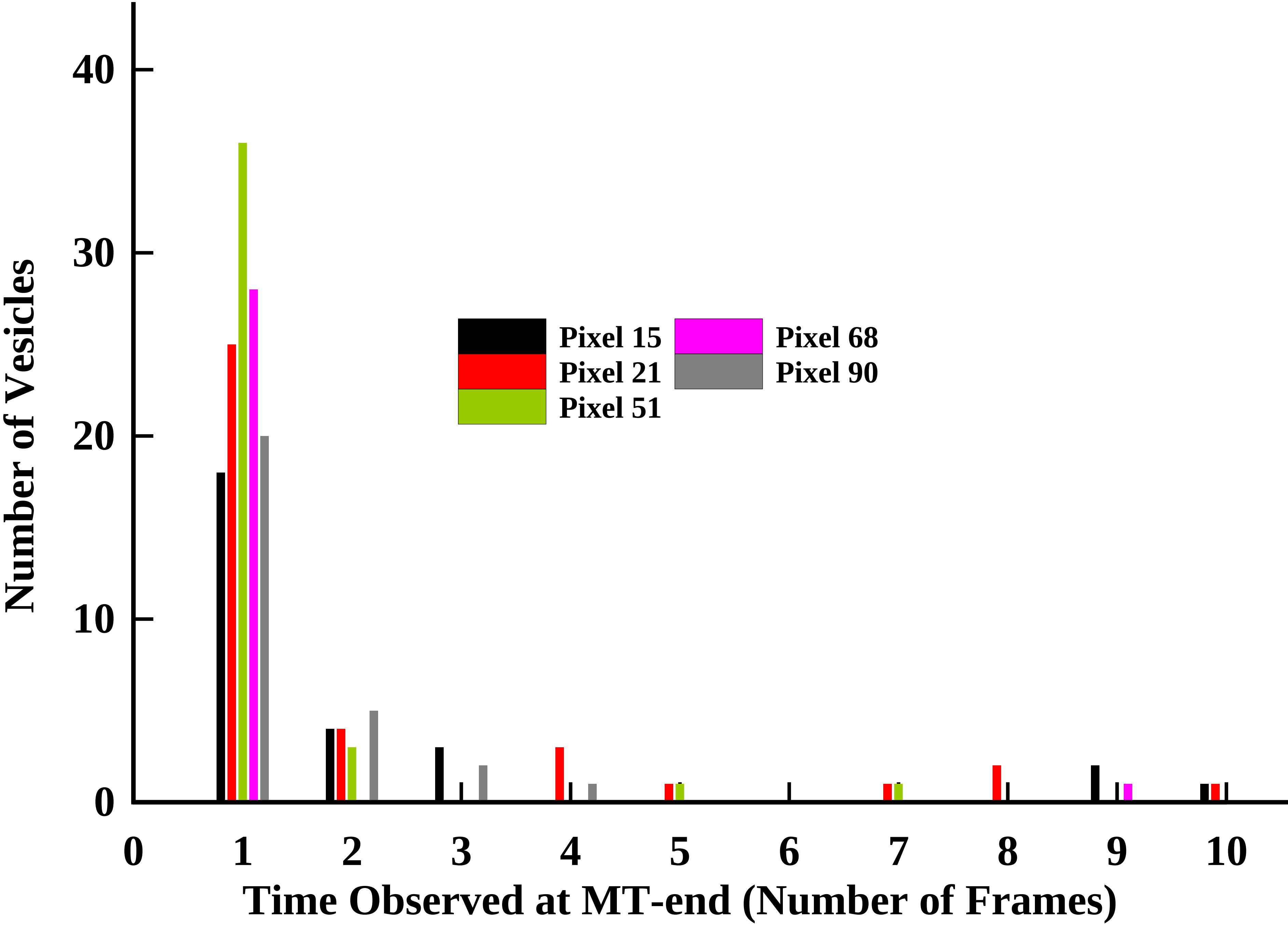

### Supplementary Figure 3

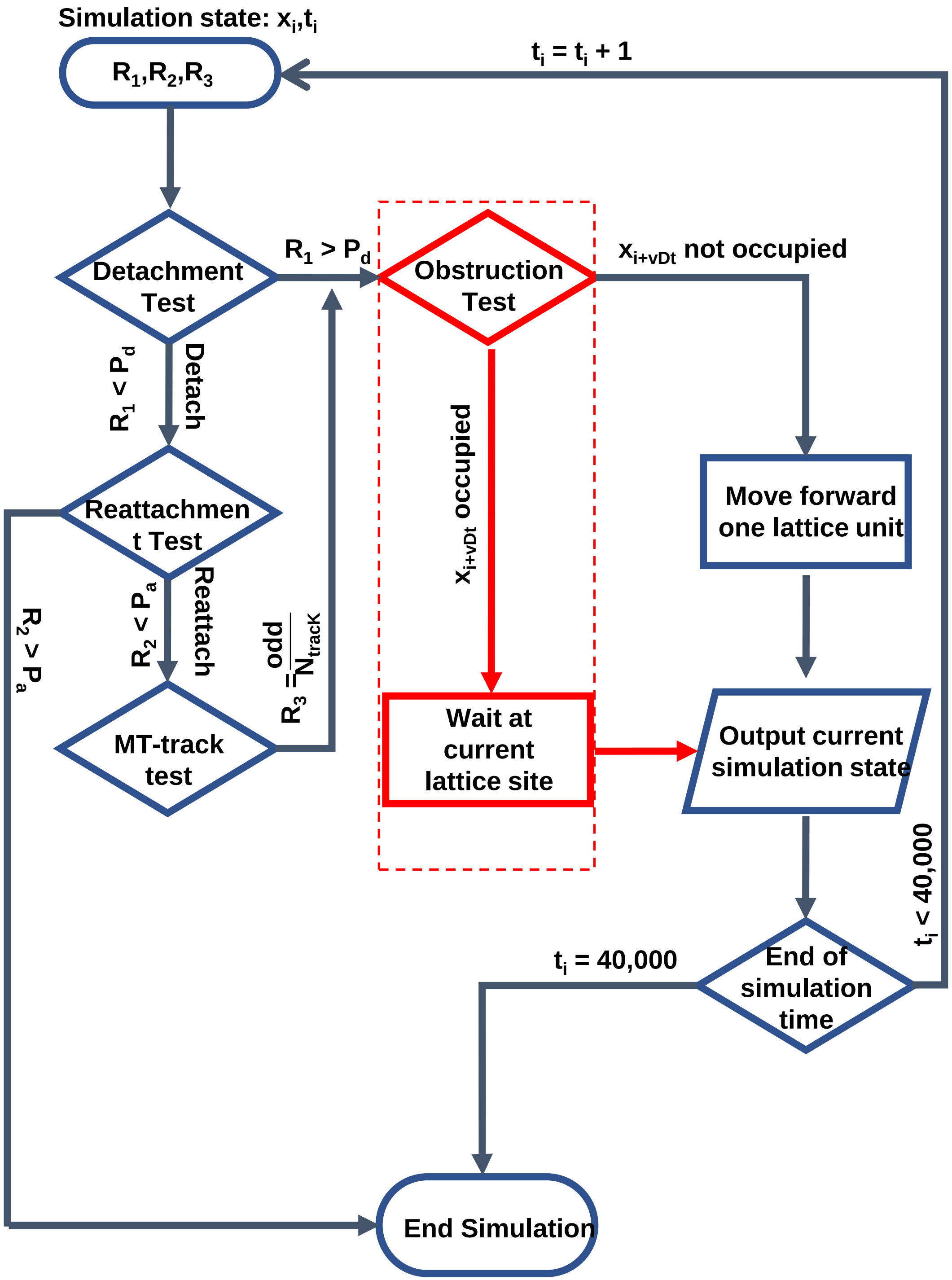

### Supplementary Figure 4

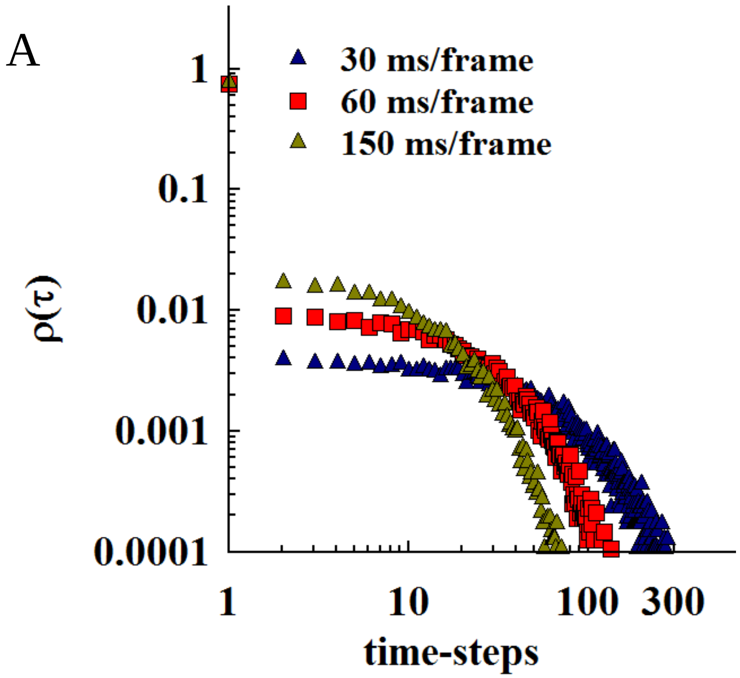

### Supplementary Figure 5

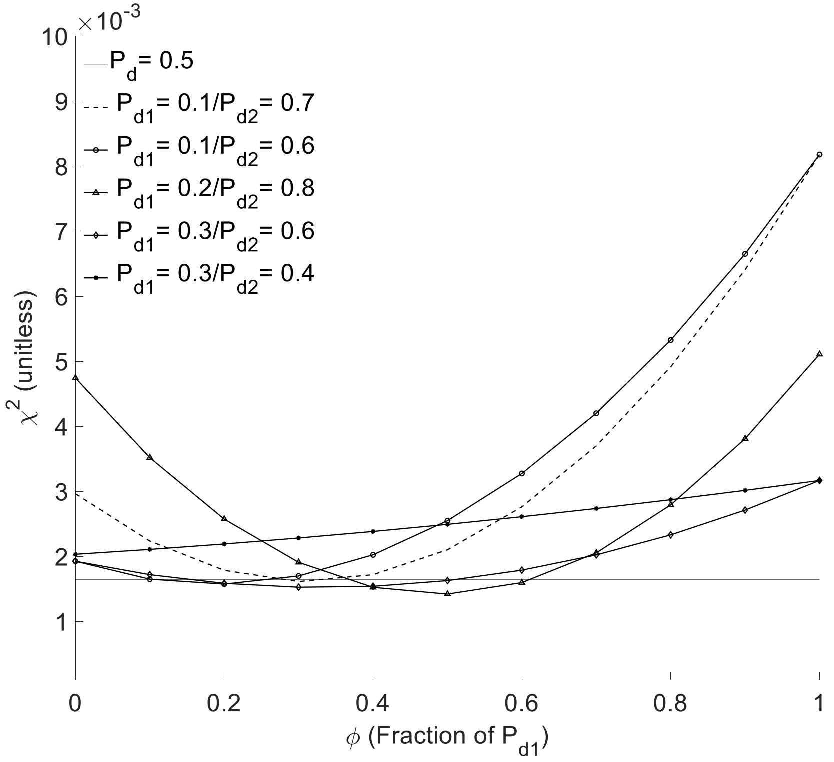

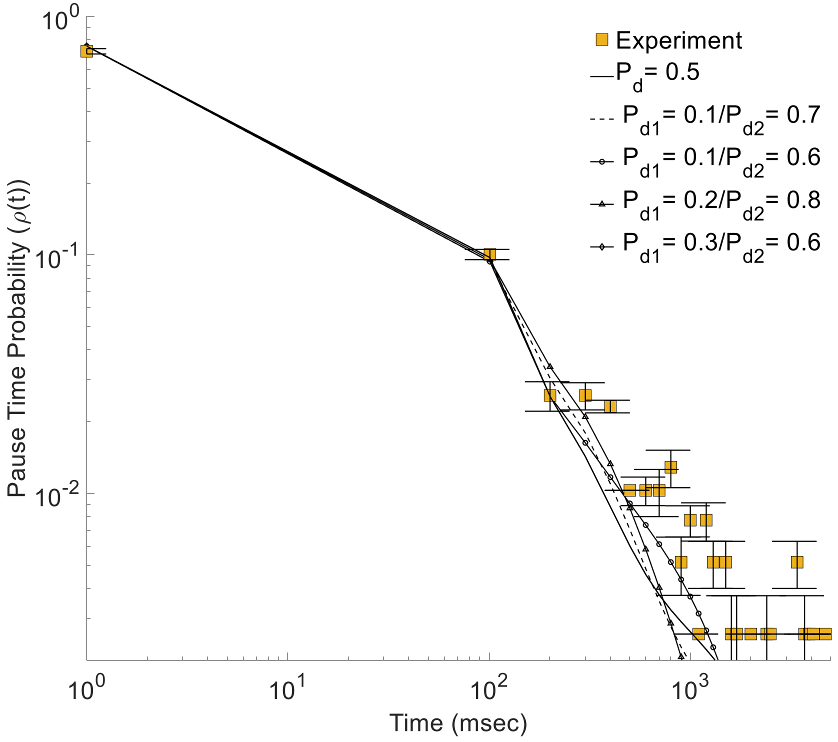
