## Supplementary Figure 2 for "Vesicle Navigation of Microtubule Ends Distinguished by A Single Rate-Constant Model"

### (A) $\tau$ = 1: Motor testing MT-end

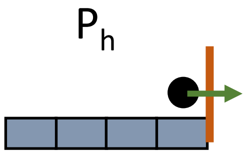


The first time a motor is inhibited by an obstruction it waits with probability **($P_h$)**

### (B) $\tau$ > [1,i] : Motor waiting at MT-end

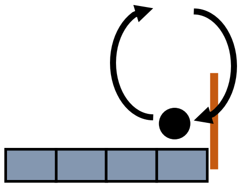

(i) Don't detach, and simply continue to wait **($1-P_d$)**

(ii) Detach, Re-attach, simply re-test obstruction **$P_d*P_h$**

### (C) $\tau$ = i+1: Motor leaves MT-end

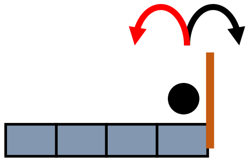

Include leaving in the next time-step **$P_d* (1-P_h)$**
