## Supplementary Table 1 for "Vesicle Navigation of Microtubule Ends Distinguished by A Single Rate-Constant Model"

| Kymograph | 30619#1b | 30619#1b | 30619#1b | 30619#1b | 30619#1b | Aggregated Data | | Distribution Data | |
|---|---|---|---|---|---|---|---|---|---|
| Pixel | 15 | 21 | 51 | 68 | 90 | | | | |
| Total-N | 28 | 37 | 41 | 29 | 28 | Total Tracks | 163 | | |
| time | | | | | | time | Aggregate | time | Fraction |
| 1 | 18 | 25 | 36 | 28 | 20 | 1 | 127 | 1 | 0.779141 |
| 2 | 4 | 4 | 3 | 0 | 5 | 2 | 16 | 2 | 0.09816 |
| 3 | 3 | 0 | 0 | 0 | 2 | 3 | 5 | 3 | 0.030675 |
| 4 | 0 | 3 | 0 | 0 | 1 | 4 | 4 | 4 | 0.02454 |
| 5 | 0 | 1 | 1 | 0 | 0 | 5 | 2 | 5 | 0.01227 |
| 6 | 0 | 0 | 0 | 0 | 0 | 6 | 0 | 6 | 0 |
| 7 | 0 | 1 | 1 | 0 | 0 | 7 | 2 | 7 | 0.01227 |
| 8 | 0 | 2 | 0 | 0 | 0 | 8 | 2 | 8 | 0.01227 |
| 9 | 2 | 0 | 0 | 1 | 0 | 9 | 3 | 9 | 0.018405 |
| 10 | 1 | 1 | 0 | 0 | 0 | 10 | 2 | 10 | 0.01227 |
